# A biomimetic deep encoder for auditory restoration by cortical stimulation

**DOI:** 10.64898/2026.11.05.764596

**Authors:** Charly Lamothe, Antonin Verdier, Corentin Caris, Ryan Greer, Romain Oliveira, Eya Mabrouk Kakaouia, Sebastian Ceballo, Pierre Platel, Emma Butt, Ella Callas, Robin San Roman, Keith Mathieson, Romain Serizel, Brice Bathellier

## Abstract

Cochlear and auditory brainstem implants are the primary treatments for profound deafness^1^. However, the inherently low resolution of electrical stimulation used by these devices limits information transfer to a small number of independent stimulation channels^2^ and results in perceptual difficulties. An alternative approach is to target the auditory cortex, a brain region large enough to accommodate hundreds of stimulation channels and whose stimulation evokes sound percepts^3,4^. Although modern brain interfaces offer large channel counts and spatial resolution, developing encoding algorithms capable of accurately transmitting interpretable sound information remains challenging. Here we introduce Braincodec, a deep network that optimizes both sound information transmission and the alignment of stimulation patterns with the spatial, temporal, and high-level properties of the auditory cortex neural code. Braincodec operates online and requires only a few hundred channels to transmit sufficient information for the reconstruction of natural sounds with high fidelity, a performance inaccessible to cochlear implant encoders. Moreover, mice equipped with a Braincodec-driven auditory cortical prosthesis matched the accuracy of non-implanted mice in discriminating real sounds. Braincodec therefore enables the development of high-throughput cortical neuroprosthetics with the potential to overcome the limitations of current auditory implants and provides a generic framework applicable to other sensory modalities.

## Main text

Multichannel neurostimulation is a key remediation strategy for severe sensory deficits such as deafness^1,2,5–7^, blindness^8–11^, or the absence of tactile inputs^12^. The prime example is the cochlear implant^2^, which provides auditory information by stimulating the auditory nerve in the ear with a small number of electrodes. Since their invention forty years ago, cochlear implants have become widely adopted^5^, as they can restore speech comprehension in quiet environments, and relatively detailed sound perception in many patients. However, cochlear implants have two fundamental limitations. Firstly, they do not address patients with bilateral auditory nerve injuries, who must instead resort to brainstem implants, providing modest benefits^7,13^. Secondly, they only provide a small fraction of the acoustic information, sufficient to understand speech in a quiet environment, but too restricted to communicate in a noisy background or to perceive the details of complex sounds like music^2^. Information loss in cochlear and brainstem implants is mainly due to poor spatial resolution of electrical stimulation in the cochlea, which results in limited distinction of different sound frequencies^2,7,13,14^. To solve these limitations, a promising strategy is to target the largest structure in the auditory system, the auditory cortex, which offers a surface area of several square centimeters, more than 10-fold larger than the cochlea or the auditory brainstem. Intraoperative electrical stimulation of the primary auditory cortex is well-known to generate clear and robust auditory perceptions^3,4^. Moreover, modern stimulation devices with submillimetric resolution, including penetrating^10,12,15^ or surface^16,17^ electrode arrays, or microLED (µLED) arrays for optogenetics^18^, enable precise stimulation of the auditory cortex with hundreds to thousands of electrodes, surpassing the 10-20 independent electrode limit of cochlear implants. However, a major missing link for developing an auditory cortical implant is an encoding algorithm tailored to the neural code for sounds in the cortex, which widely differs from the neural code at the auditory system periphery. The cochlea decomposes sounds along a one-dimensional tonotopic axis with high temporal precision on the order of milliseconds^19^. The primary auditory cortex encodes sounds at a >10 fold lower temporal resolution^20,21^ across a two-dimensional tonotopic map^22,23^. Moreover, the auditory cortex integrates sound information over longer timescales^24,25^, displays complex offset dynamics at sound termination^26,27^, and employs diverse rate-based codes for a broad range of acoustic features^28–31^. No computational solution could thus far reproduce these features while ensuring the full preservation of auditory information as needed for boosting auditory restoration quality. Here, we used training objectives known to generate sound representations in deep neural networks similar to those observed in the auditory cortex^28,32,33^ and combined them with an optimized framework for high-performance online sound compression^34^. This led to Braincodec, an algorithm that provides biomimetic stimulation patterns for a cortical implant while minimizing information loss. By reconstructing sounds from the stimulation patterns with an optimized decoder, and comparing reconstructed to original sounds, we show that Braincodec transmits enough information, with only 256 channels, to restore all perceptually relevant acoustic details of natural sounds, including sound source localization from binaural auditory inputs. By contrast, cochlear implant encoders lose a large part of this information. Moreover, mice equipped with a Braincodec-driven cortical implant prototype experience perceptions that they could discriminate as precisely as real sounds in a behavioral task. Braincodec therefore paves the way for a new generation of neurostimulation devices, which could reach a level of auditory restoration close to natural hearing.

### Efficient sound compression

To design Braincodec, we reasoned that a sound-encoding algorithm for a neuroprosthetic device should simultaneously solve two objectives by: (1) providing all the information necessary to reconstruct the original sounds, which is possible based on auditory cortex activity^35,36^, and (2) respecting the spatial and temporal organization of the neural code. To solve these two objectives, we trained a deep neural encoder together with a deep neural decoder which reconstructs encoded sounds to minimise information loss, and a classifier that steers the stimulations patterns towards biomimetic neural codes (**Fig. 1a,b**). The Braincodec deep neural encoder has 14-layers. It converts the sound waveform sampled at frequency FS into spatio-temporal stimulation patterns for N channels at frequency FC << FS (**Fig. 1a**). We chose FC = 40 Hz, based on the highest amplitude modulation frequency followed by most cortical neurons and on estimates of the boundary between pitch and flutter for amplitude-modulated sounds^20,37^. The conversion is performed by one-dimensional causal convolutional filters allowing low-latency, online audio processing^34,38^. The deep neural decoder mirrors the encoder architecture. It is trained to reconstruct, as accurately as possible, the original sound based on the encoder output (**Fig. 1b**). The perceptual quality of the sound reconstruction is optimized with a multiscale adversarial strategy that discards spurious features in the generated sounds^34^. The stimulation patterns generated by the encoder under the control of the decoder can be seen as the latent representation of sounds in an autoencoder network^39^, which minimizes information loss (**Fig. 1a, b**). This architecture is appropriate to solve Braincodec’s information preservation objective, but does not resolve the biomimetic encoding objective, because, generally, autoencoders poorly reproduce biological representations observed in sensory cortex^28,33,40^. Sensory systems in the brain not only transfer accurate information, they also contribute to the recognition of objects and behaviorally relevant stimulus categories. This is likely the reason why deep networks which are able to solve such recognition tasks generate representations highly similar to sensory cortex activity, both in hearing and vision^28,32,33,41^. Therefore, to fulfill the second objective of the encoding algorithm, we added a shallow classifier network that uses the encoder output to identify 423 sound classes, corresponding to broad categories of sounds (e.g. Mechanisms, Music, Wind, Conversation, Surface contact **Fig. 1b**).

**Fig 1:**
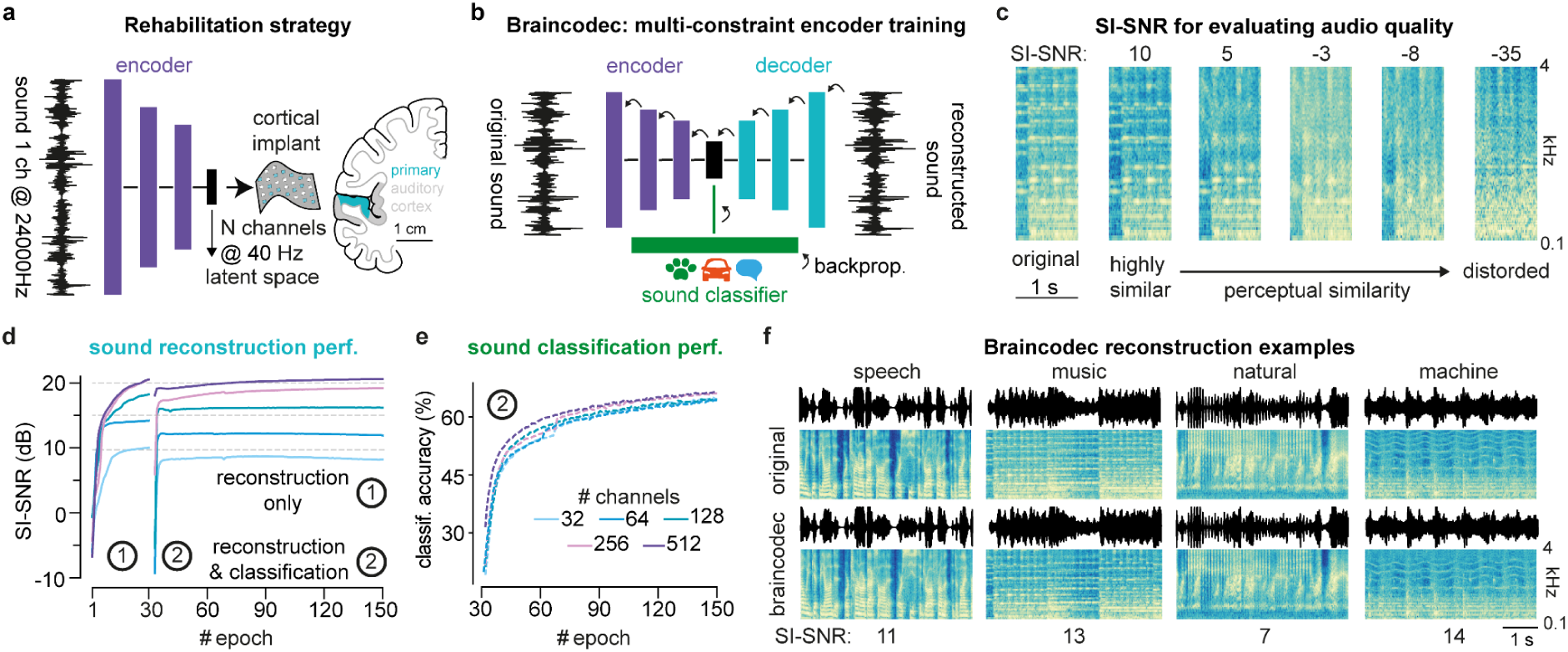
Braincodec efficiently compresses sound information. **a**. Schematic of the auditory cortical implant. The purple bars represent the layered architecture of the deep encoder network. **b**. Architecture of the networks used to train the Braincodec deep encoder (purple). The blue and green bars represent the layered architecture of the decoder and classifier networks, respectively. The arrows indicate the flow of error backpropagation (backprop.). **c**. Spectrogram of a 1 s music sample and of the same sample reconstructed with decreasing accuracy to illustrate the correspondence between sound degradation and SI-SNR value, shown above the spectrograms. Frequency scale is logarithmic. Blue and yellow color indicate low and high magnitude respectively. **d**. Mean SI-SNR of reconstructed sounds (validation dataset) during the two-phase training of Braincodec for 5 different channel counts. In one training epoch, 5796000 sound snippets are reconstructed and classified. Phase ①: reconstruction training only. Phase ②: reconstruction and classification training in parallel. **e**. Classification accuracy during training phase ②. Perf. = performance. **f**. Spectrogram of sample sounds and of their reconstructions with the 512 channel Braincodec algorithm (decoder + classifier).

We trained Braincodec to compress, reconstruct, and classify 5.8 million human-labeled natural sounds of 2 s duration. We assessed the similarity of the reconstructed and original sounds using the scale-invariant signal-to-noise ratio metric^42^ (SI-SNR). A SI-SNR below 0 reflects poor similarity between the original and reconstructed sounds, 5 corresponds to a good similarity, 10 means that the two sounds are hard to distinguish (**Fig. 1c)**. This scale is illustrated by the audio samples provided in **Supplementary Video 1**. After training, Braincodec could reconstruct sounds at an average SI-SNR above 0 with only 64 channels and reached close to 10 with 256 or 512 channels (**Fig. 1d**), independent of the sound type (**Extended data Fig. 1**), while performing sound classification above 60% accuracy almost independently of channel count (chance level: 0.2%, **Fig. 1e**). This demonstrates that highly-efficient encoding of any sound onto only 256 or 512 channels is possible at the low temporal resolution of the auditory cortex, as also illustrated by the spectrograms, waveforms, or audio samples of reconstructed and original sounds (**Fig. 1f**, **Supplementary Video 2**). Moreover, information transmission performance is not compromised by the parallel resolution of a difficult sound classification task.

### Neuromimetic sound representations

To assess the similarity between Braincodec’s output and sound-driven activity in the auditory cortex, we first used a publicly-available dataset of human cortex sound responses recorded with functional magnetic resonance imaging (fMRI)^43^ (**Fig. 2a**). Focusing on the primary auditory cortex, we computed the linear function that best predicts fMRI responses based on the Braincodec encoder output for a ‘training’ subset of sound responses. We then used the percentage of explained variance in the observed neural responses for a ‘test’ subset of sounds as a metric of similarity between Braincodec and neural data (**Fig. 2b**). Based on this procedure, Braincodec representations could explain up to 71% of auditory cortex activity (**Fig. 2c-d**). This level of performance is close to state-of-the-art deep networks for sound recognition^32,33^ which do not preserve acoustic information (**Extended Data Fig. 2a-d**). By contrast, the compression-only autoencoder networks explained no more than 19% of observed sound responses (**Fig. 2c-d**). The similarity between Braincodec and brain activity was not affected by the sound reconstruction task and was high for all tested channel counts, peaking at 256 channels (**Fig. 2d**). This highlights the compatibility of the classification and reconstruction objectives, and the importance of the classifier module in Braincodec for the generation of a biomimetic code. Similarly, it was not possible to reconstruct sounds accurately from a classifier trained without the decoder (**Extended Data Fig. 2b-d**). Hence, the association of a sound classification and sound reconstruction task in the Braincodec training procedure provides a unique sound encoding solution, preserving almost all acoustic information in a code resembling auditory cortex activity.

**Fig 2:**
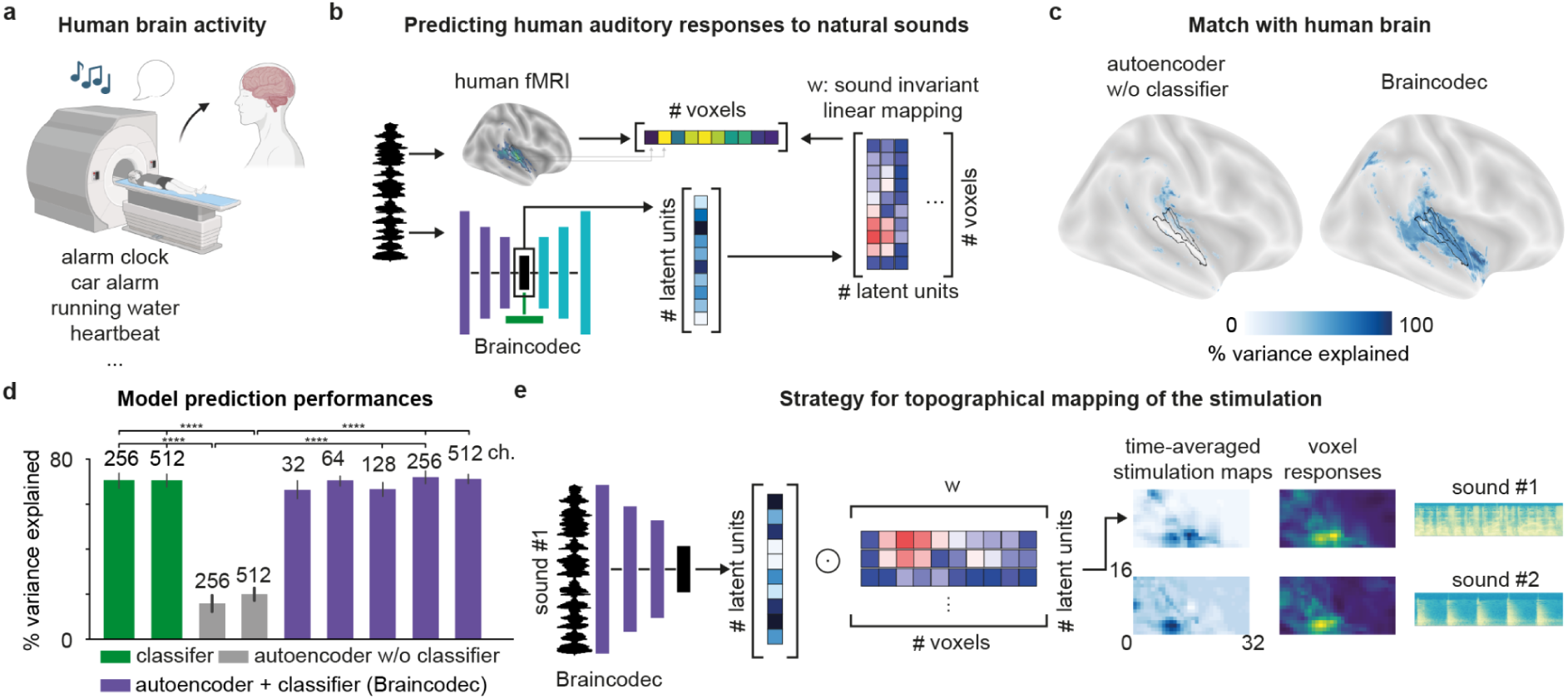
Braincodec reproduces human sound representations. **a**. Illustration of the fMRI-based brain activity recordings. **b**. Linear mapping of Braincodec representations to fMRI voxels to estimate representation similarity. **c**. Fraction of variance explained by the autoencoder network alone and by the Braincodec encoder for voxels localized in the temporal region of the brain. **d**. Fraction of explained variance for Braincodec with different channel counts, and for the autoencoder or classifier alone. (**** p=3.0518 x 10⁻⁵, Bonferroni corrected paired Wilcoxon signed-rank tests, n = 16 (median A1 voxels from 8 subjects * 2 hemispheres). **e**. The linear mapping procedure to fMRI recordings allows mapping Braincodec outputs to the physical electrodes and their target location of the auditory cortex.

The comparison of Braincodec and fMRI recordings of auditory cortex activity also provide a methodology to align the cortical implant output to the spatial organization of the auditory cortex in every human subject. Our analysis (**Fig. 2c-d**) shows that once computed on a set of functional responses to a natural sound sample, the linear function which maps Braincodec’s output to auditory cortex activity is effective for any natural sound and can be used as the spatial alignment function (**Fig. 2e**). While functional maps cannot be derived in profoundly deaf patients, anatomical specificities of a given patient could be estimated based on response maps in anatomically-similar hearing subjects. This provides a concrete solution to position the cortical implant on the patient’s brain at the mesoscopic scale, bearing in mind that electrical stimulation of the cortex will only coarsely approximate the functional organization of the auditory cortex at the scale of single neurons which display substantial variations across neighboring neurons^44^.

### Temporal dynamics and feature tuning

fMRI recordings give access to the spatial representation of sounds at the scale of the entire human auditory cortex but have limited spatial and temporal resolution. If Braincodec single-channel activity is examined for a range of simple and more complicated sounds, complex spatio-temporal dynamics are observed (**Fig. 3a-b**). These resemble electrophysiological neural responses to sounds observed in the auditory cortex of animals or humans. For example, some Braincodec channels activate transiently at the onset or offset of pure tones while other channels activate in a sustained manner (see sample traces in **Fig. 3b** and quantification in **Extended Data Fig. 3**). These three types of dynamics are commonly observed in the auditory cortex^26,30,45,46^. As in the cortex^26,46^, sustained, onset and offset activations could be the only component of the response or could be combined in the same response (**Fig. 3a-b**, **Extended Data Fig. 3**). Furthermore, offset responses or isolated onset responses were seldom observed in the autoencoder network lacking the classifier used in Braincodec (**Fig. 3a-b**; fractions of all pure tone responses with offset activation: 0.4% autoencoder only, 9.4% Braincodec; fractions of all pure tone responses with onset activation only: 0.9% autoencoder only, 8.8% Braincodec; **Extended Data Fig. 3**). This observation suggests that transient responses, and particularly offset responses, reflect computations that enable the robust recognition of meaningful sound classes.

**Fig 3:**
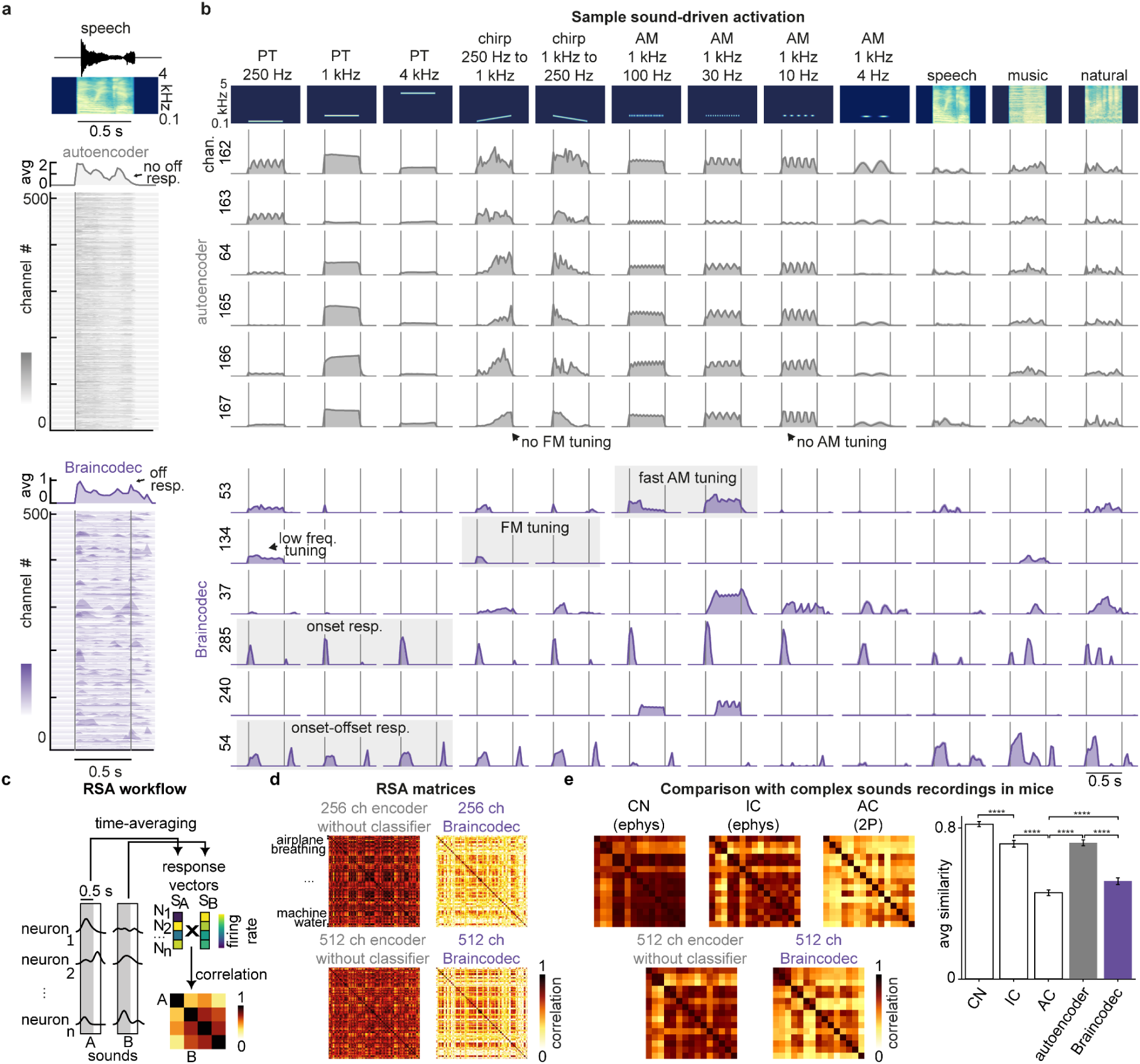
Braincodec resembles biological sound response dynamics. **a**. Response time courses for the 512 channels of the autoencoder without classifier, and of Braincodec, for a sample speech sound. The color intensity (see colorbar) represents channel activity level (0 to 50 arbitrary units). **b**. Activity of sample channels from the autoencoder without classifier and from Braincodec (512 channels) for pure tones (PT), frequency-modulated (FM) chirps ramping up or down, amplitude-modulated (AM) tones at 4 frequencies and 3 natural sounds. Grey shadings highlight feature-tuning behavior similar to real auditory cortex neurons. **c**. Construction of the representation similarity analysis (RSA) matrix which measures similarities of representations across a set of sounds. **d**. RSA matrices for a set of natural sounds encoded by the autoencoder without classifier and by Braincodec. **e.** (top row) noise-corrected RSA matrices for neural data recorded in the mouse cochlear nucleus (CN, 1421 neurons), inferior colliculus (IC, 551 neurons), and auditory cortex (AC, 4217 neurons) by Gosselin et al., 2025^31^, corresponding to responses to 15 complex natural sounds. (bottom row) RSA matrices for the representations of the same sound by the 512 channel autoencoder trained without classifier and the 512 channel Braincodec. Latent codes were averaged over time before computing correlations across sounds. (right) Mean pairwise similarity across the 105 unique sound pairs (± SEM). Representation similarity decreased along the auditory pathway (CN > IC > AC). Statistical comparisons used two-sided Wilcoxon signed-rank tests on the 105 paired pairwise similarities: CN vs IC, W = 491, p = 2.4 × 10⁻¹³; IC vs AC, W = 51, p = 2.5 × 10⁻¹⁸; AC vs autoencoder, W = 17, p = 9.5 × 10⁻¹⁹; autoencoder vs Braincodec, W = 272, p = 1.0 × 10⁻¹⁵; AC vs Braincodec, W = 1456, p = 2.2 × 10⁻⁵.

Braincodec output activations also display prominent tuning to stimulus features beyond frequency selectivity as commonly observed in single neurons of the auditory cortex^30,31^. This included tuning to temporal features such as the direction of frequency modulations (**Fig. 3b**)^47–49^ or the frequency of amplitude modulations^20,21^ (**Fig. 3b, Supplementary Audio 1, Extended Data Fig. 3**). The progressive emergence of feature tuning in the auditory system reflects a more general decorrelation process, through which the time-averaged representations of most sounds become less similar to each other, as quantified recently in the mouse using representation similarity analysis (RSA)^28,31^. In line with this, natural sound representations in Braincodec’s output were less correlated than in the autoencoder network alone (**Fig. 3c-e**). Moreover, the average correlation between sound representations and the structure of the RSA matrix for Braincodec were similar to previous measures from the auditory cortex and dissimilar to measures from subcortical auditory centers^31^ (**Fig. 3e**). Together, Braincodec produces sound representations that share major properties with those observed in the auditory cortex (**Figs. 2&3**), while preserving an excellent information throughput (**Fig. 1**), making it a highly promising solution to perform sound encoding in a cortical implant for auditory rehabilitation.

### Braincodec outperforms cochlear implant encoders

To estimate the potential benefits of a Braincodec-operated cortical implant against cochlear implants, we directly compared their theoretical information throughputs. The sound encoder in cochlear implants measures the energy contained in several frequency bands computed from the Fourier Transform of the sound over a short time window (**Fig. 4a**) to determine the current amplitudes sent through the stimulation channels^2^. Some cochlear implant encoders select the N channels among a total of M that have the highest energy (typically M = 22, N = 12) and zero all other channels^50^; other encoders skip this step, usually in implants with lower channel counts. The energy values are then passed through a logarithmic compression function to compute the current amplitudes applied in each of the N active channels (**Fig. 4a**). Several approaches have been used to estimate the acoustic information transmitted, including linear vocoders based on the frequency band energy before channel selection^51^, or neural vocoders^52^. Here, to provide a fair comparison with Braincodec, we trained 1D convolutional deep neural network decoders with the same architecture as Braincodec’s decoder to maximize similarity between the original sound waveform and its reconstruction based on different encoding steps for the cochlear implant (**Fig. 4a**).

**Fig 4:**
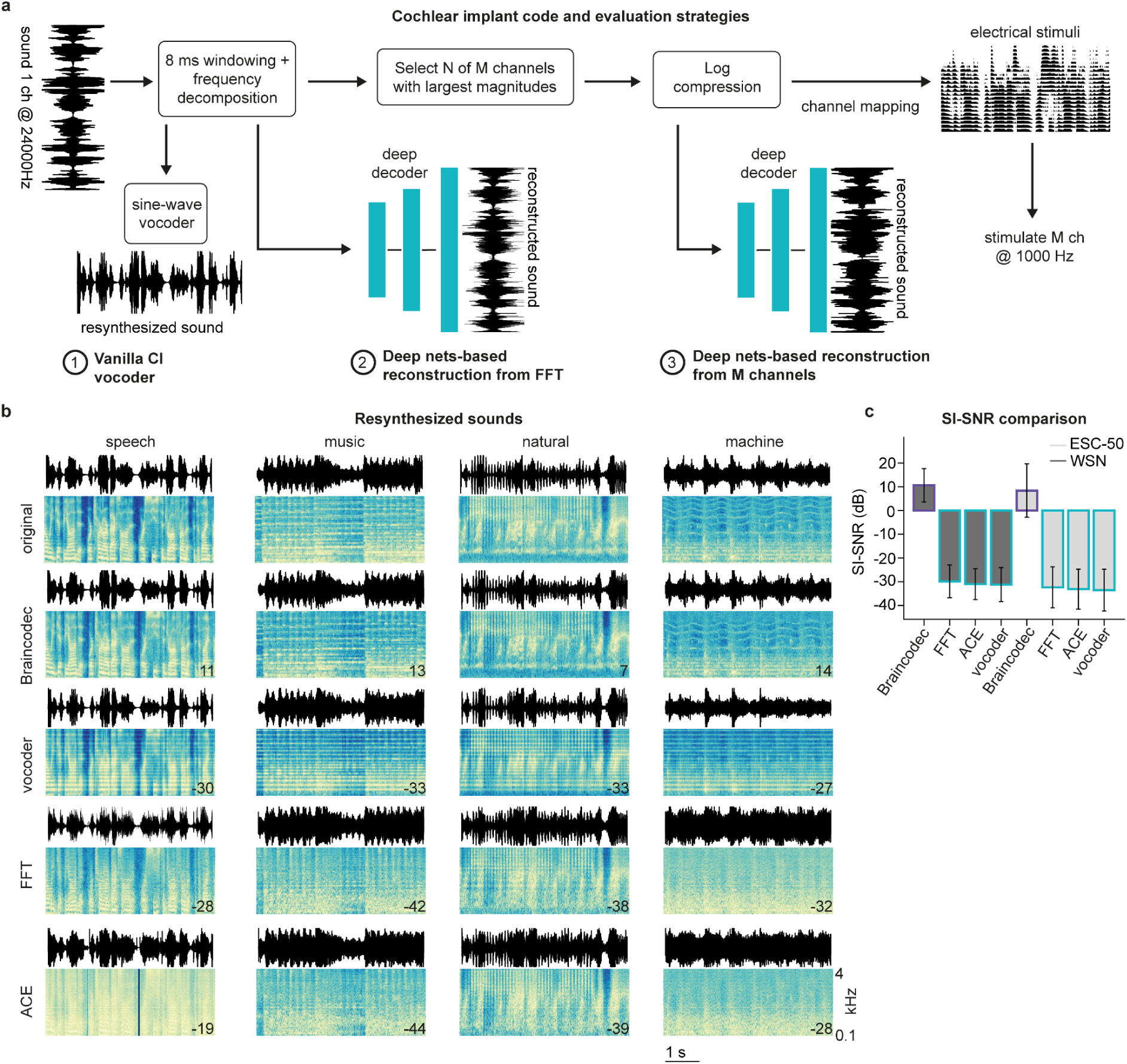
Braincodec outperforms cochlear implant encoders. **a**. Block diagram of the common cochlear implant speech processor algorithm, advanced combination encoder (ACE), and of the three sound reconstruction strategies used to evaluate the information loss of the ACE sound encoder using a linear vocoder for 22 channels (① vocoder), a deep network-based approach after the 22 channel FFT step (② FFT) or after the final step of ACE (③ ACE) **b**. Spectrograms and sound waves for different sounds and their reconstruction by Braincodec with 512 channels and the 3 cochlear implant reconstruction strategies depicted in **a**. The number in the bottom right corner of the reconstructed spectrogram is the SI-SNR value indicating reconstruction accuracy (10 excellent, 0 to −10 poor, <10 very poor, see Fig. 1c). **c**. Average SI-SNR (mean ± standard deviation) of reconstructed sounds for the evaluation (WSR) and validation datasets (ESC-50). SI-SNR for Braincodec is significantly different from the SI-SNR for cochlear implant reconstructions (Braincodec vs Vocoder: p < 0.001; Braincodec vs FFT: p < 0.001; Braincodec vs ACE: p < 0.001; paired two-sided t-tests, Benjamini-Hochberg FDR corrected, n = 2000 sounds).

Using this approach, we observed that cochlear implant encoders discard a large amount of key acoustic details, consistent with previous theoretical and perceptual estimates^51^. This was indicated by average SI-SNR values around −30 (**Fig. 4b-c** *vs* around +10 for Braincodec, **Fig. 1d,f**) and poor spectrogram reconstruction (**Fig. 4b**). Qualitatively, sample sounds reconstructed with our method from cochlear implant encoders (**Supplementary Videos 3-6 and Audio 2**) allow speech understanding despite strong deterioration of the sound, but harmonies in music and acoustic fine structure are lost, in line with the experience of cochlear implant users^2^. The mean reconstruction SI-SNR after computing sound energy in 22 frequency bands is −32.3 +/- 8.6 and only slightly reduces to −33.1 +/-8.4 after N of M band selection for the ACE encoding algorithm^50^ (validation dataset, ESC-50, **Fig. 4c**). Reducing the number of channels without band selection leads to similarly poor reconstructions (**Extended Data Fig. 4, Supplementary Audio 2**). Hence, most of the information loss in the cochlear implant encoder happens at the initial encoding stage when computing the energy of frequency bands. These theoretical estimates confirm that the acoustic information throughput of a cochlear implant is largely outperformed by Braincodec with a few hundred channels (**Fig. 1d,f**), despite the lower clock frequency of Braincodec (40Hz) compared to cochlear implants (1kHz).

### Precise discrimination of Braincodec-based stimulations in mice

To evaluate the effectiveness of Braincodec patterns to elicit perceptions, we used a recently developed implantable optogenetic stimulator^18^, made of a 10 x 10 LED array (2-by-2 mm^2^ total size) (**Fig. 5a-b**), connected to a dedicated electronic system enabling precise control of arbitrary spatio-temporal patterns (**Extended Data Fig. 5**). The LED array was chronically implanted over the A1 subfield of the primary auditory cortex (**Fig. 5c-d**). Expression of the opsin ChRmine in pyramidal neurons (CamKII promoter) was obtained with transgenic mice^53^ or adeno-associated virus (AAV) injections. Braincodec is designed to process natural sounds and is effective for sounds in the 200-4000 Hz frequency range in which the relevant information and energy is concentrated^54^. The mouse auditory cortex encodes sound frequencies above ∼ 3000 Hz, corresponding to the hearing range of the species. We therefore constructed Braincodec patterns for two pure tones in the human hearing range (500 and 2000 Hz, **Fig. 5e-g**), and aligned them to the mouse tonotopic map to cover a similar frequency difference of 2 octaves (4 to 16 kHz, **Fig. 5d**, **Extended Data Fig. 6**). Such a transposition along the frequency dimension is meaningful for pure tones but would be difficult for natural sounds which we did not use in these experiments. The Braincodec patterns were generated with a 100 channel version of the algorithm. This version was trained with an additional objective function that maps pure tone patterns to the tonotopically organized responses of the mouse auditory cortex along the frequency axis of A1 (see Methods, **Extended Data Fig. 6**). The relative weight of this tonotopic constraint was tuned to effectively generate patterns that follow the tonotopic axis but are not strictly organized (**Fig. 5a**) as observed in the mouse auditory cortex^44^.

**Fig 5:**
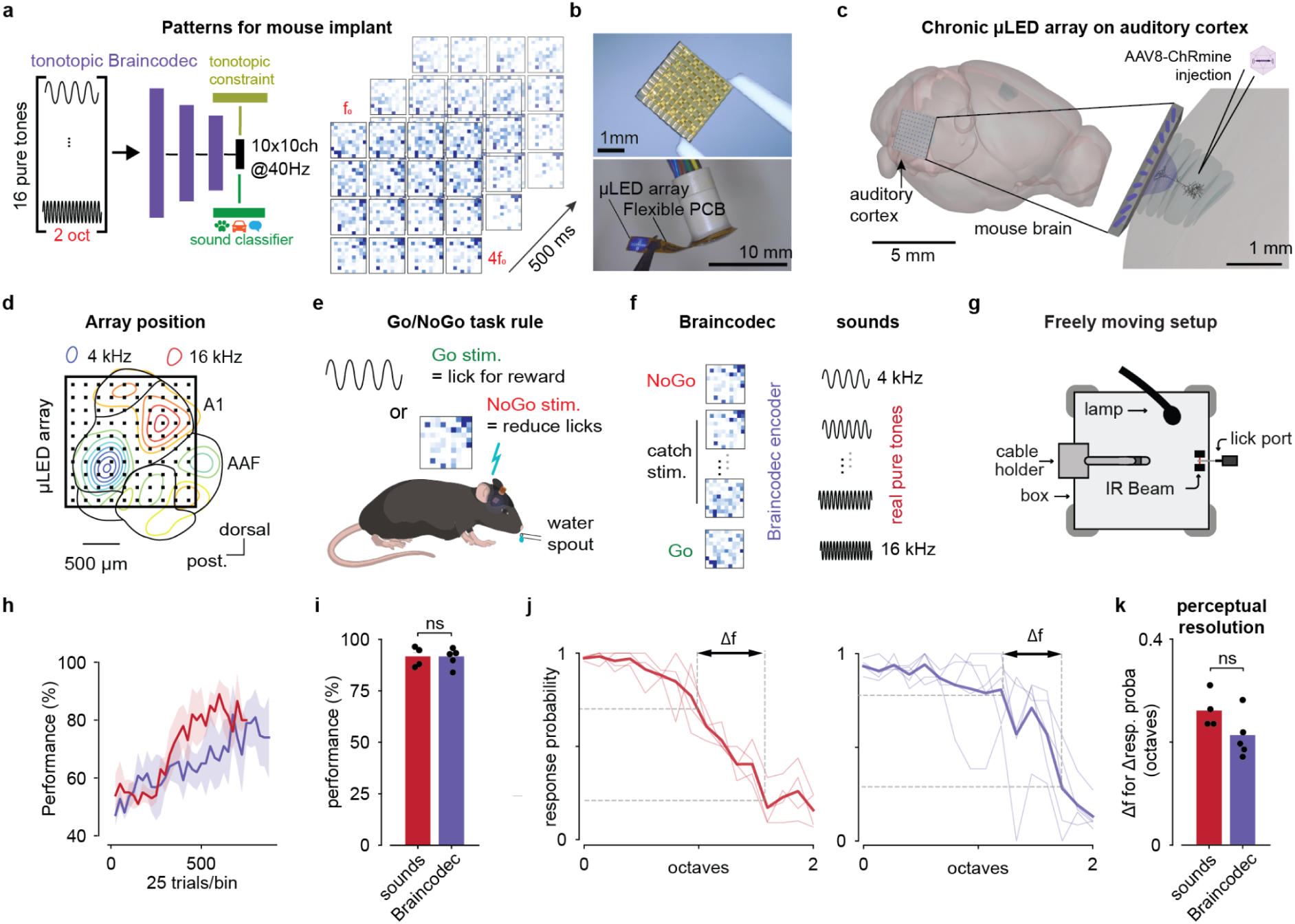
Braincodec produces precise perceptions in mice. **a**. (left) Schematic of a variant of Braincodec including a spatial mapping constraint (the decoder network is omitted from the schematic but is included in the architecture) to reproduce a single axis tonotopy. (right) Output activity for 16 pure tones over 2 octaves including 3 time points (0-25, 225-250, 475-500 ms). **b**. Pictures of the implantable µLED array. **c**. Implantation strategy. **d**. Position of the LED array on the auditory cortex targeting the main tonotopic field A1. AAF: anterior auditory field. **e**. Task rule for the Go/NoGo discrimination task. **f.** Categorization task for Braincodec patterns and pure tones discrimination. In both tasks, mice are trained to discriminate between the two extreme stimuli and are subsequently tested with catch stimuli corresponding to intermediate tone frequencies between these extreme frequencies. **g.** Sketch of the behavioral setup. **h**. Group-averaged learning curves for the Braincodec-trained (purple) and sound-trained (red) mice (grey shading = SEM; no significant group effect on learning curves, Wilcoxon ranksum test on the number of trial to 80% accuracy, p = 0.56, sounds: n = 4, Braincodec: n = 5 mice). **i**. Discrimination performance during the categorization task (two-sided Wilcoxon rank sum test; n = 4 and 5 mice; p = 0.90). **j**. Response probability during spontaneous categorization of intermediate sound (red) or Braincodec patterns (purple). Thick line: population average; thin lines: single mice; dashed lines sketch the perceptual resolution measure. **k**. Perceptual resolution measured as the minimal frequency difference for which response probabilities differ by 0.5. Two-sided Wilcoxon rank sum test, n = 4 and 5 mice; p = 0.11; dots: single mice; bar: group average.

We trained a group of implanted but freely moving mice to discriminate the two Braincodec patterns in a Go/NoGo discrimination task. As a comparison, we also trained a group of non-implanted mice to discriminate between two real pure tones corresponding to the frequency locations where low and high frequency Braincodec patterns were targeted (4 and 16 kHz, **Fig. 5e-g**). Both mouse groups displayed similar learning curves (**Fig. 5h**), showing that Braincodec patterns are discriminable by mice despite their complexity and their coarse alignment to the high-resolution spatial organization of sound representations^44^.

To estimate the accuracy of the resulting perception, we measured how mice categorize, in rare catch trials, 14 Braincodec patterns corresponding to 14 pure tone frequencies logarithmically-spaced between the two target frequencies of the Go/NoGo task (**Fig. 5f**). The same test was performed with real pure tones in the second mouse group (**Fig. 5f**). Both groups displayed the same level of correct responses for the Go and NoGo sounds during this categorization assay (**Fig. 5i**). To estimate perception accuracy, we measured the smallest relative frequency difference (Δf) leading to a difference of 0.5 response probability (**Fig. 5j**). Pure tone categorization led to similar and non-significantly different Δf values between mice using Braincodec information and mice using pure tones (pure tones : 0.26 +/-0.02 octave; Braincodec : 0.21 +/- 0.02 octave, mean +/- SEM, **Fig. 5k**). This demonstrates that the resolution of perceptions generated by Braincodec patterns is similar to the resolution obtained with real pure tones in mice. To compare the performance of Braincodec to a potentially simpler approach in which sound frequency is encoded by an idealized tonotopic code in strict frequency bands, we trained another group of mice to discriminate, under head-fixation, thin optogenetic stimulation bands delivered with a precise videoprojector system and placed perpendicularly to the tonotopic axis (**Extended Data Fig. 7a-f**). While head-fixation had little effect on the precision of discrimination for pure tones, Δf values for this idealized tonotopic stimulation approach were two times larger than the Δf values observed for Braincodec stimulation patterns (**Extended Data Fig. 7g-i**). This confirms that the two-dimensional patterns of Braincodec are effective in transmitting resolved information to the auditory cortex, which can be used by mice to perform discriminative behavior.

### Binaural processing with minimal performance loss

Another advantage of a cortical implant is that the auditory cortex of each cortical hemisphere receives and processes information from the two ears, despite a dominance of the contralateral spatial hemifield in the primary auditory cortex^3,55^. Spatialization of sound perception is needed for proper hearing restoration and bilaterally implanted cochlear implant users report greater comfort despite the limited transfer of spatial cues by the devices. We therefore tested whether it was possible to add binaural spatial information to Braincodec. We modified the encoder to receive two sound waveforms corresponding to acoustic inputs at the left and right ears, simulated with experimentally-measured head-transfer functions^56^ for a range of locations in a grid covering 75% of the frontal hemifield (**Fig. 6a**). This spatialized Braincodec was then trained to optimize three objectives: reconstruct the two sound inputs, classify sound identities and regress sound direction. The sound direction regression was performed by extending the classifier network with a regression module (**Fig. 6b**). Training with these three objectives was successful for both 256 and 512 channels. Sound classification and similarity of Braincodec stimulations to human brain activity were minimally affected by the spatial information processing (**Fig. 6c-d**). At the same time, spatial location regression was largely above chance (**Fig. 6c**). In addition, sound reconstruction accuracy only dropped from a SI-SNR of ∼9 for 256 channels and ∼10 for 512 channels (**Fig. 1a**) to 5 and 7 respectively (**Fig. 6c, Supplementary Audio 3**). This maintains good information transmission quality and is consistently better than the cochlear implant encoders (**Fig. 4**). The small drop of reconstruction performance further indicates that the representations necessary to solve the double objective of maintaining acoustic information, and providing high level interpretations of the sound, are largely compatible. Importantly, the binaural version of Braincodec produced channel activations with spatial tuning properties largely observed in the mammalian auditory cortex^55,57^. Some channels were most active for sounds arriving from the left, the right, or the center of the simulated environment (**Fig. 6e**). In some cases, sound direction tuning could be modulated by sound identity in the same neurons (**Fig. 6e**). Hence, biologically-realistic features of the code for spatial cues in sounds can be introduced in our cortical implant encoding algorithm, with a limited loss of information about other sound features.

**Fig 6:**
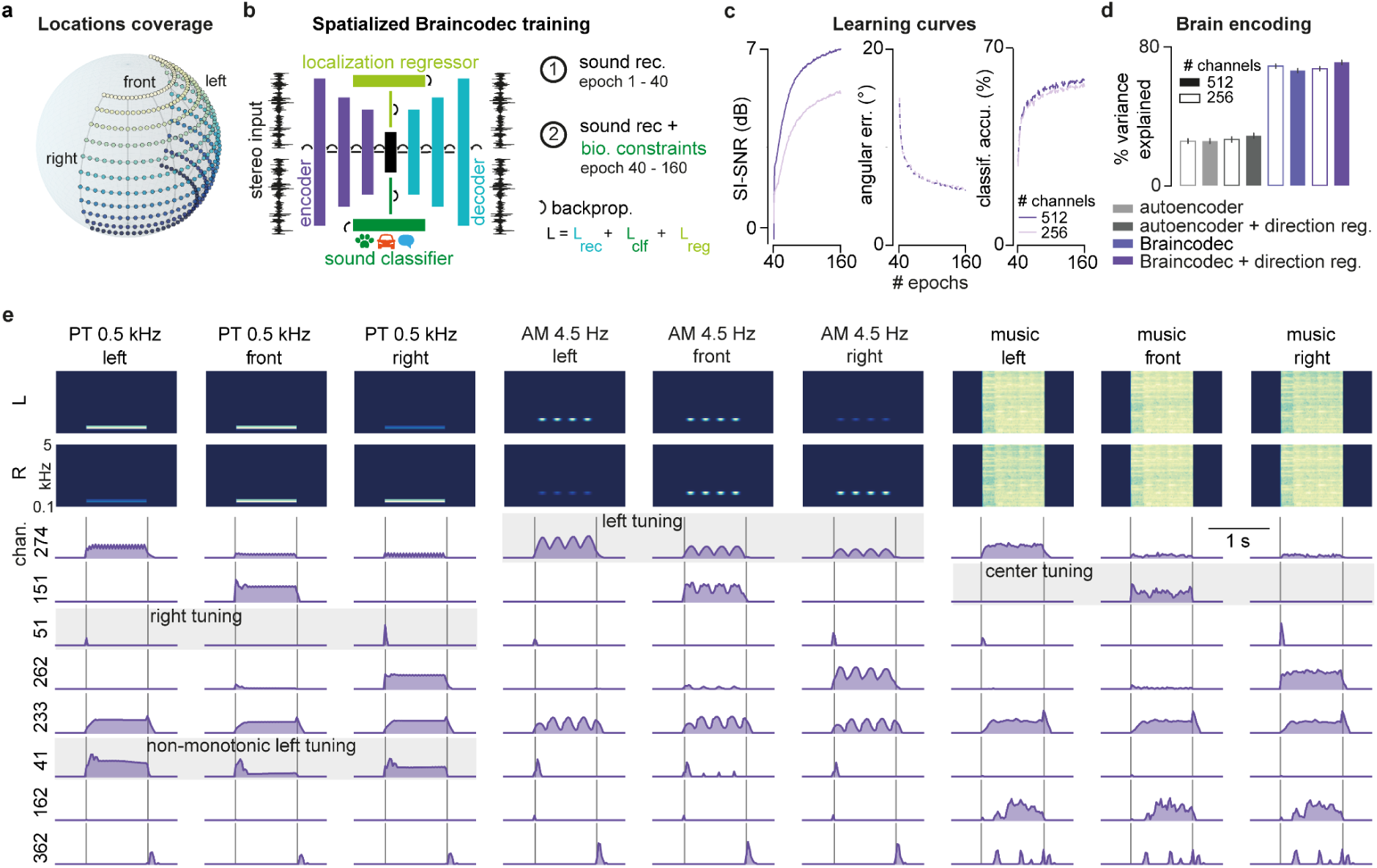
Binaural processing induces minimal performance loss. **a**. Simulated sound direction grid. **b**. Schematic of the Braincodec architecture for processing spatialized binaural sound inputs. The network processes two sound channels and training includes a regression of sound direction. **c.** Learning curves for the reconstruction (SI-SNR, solid line), sound identity classification (acc., long dashes) and direction regression (loc., short dashes) tasks **d**. Fraction of explained variance in human auditory data (same as Fig. 2) for Braincodec and for the autoencoder or classifier alone; trained with and without the sound direction regression task. **e**. Activity of sample channels from binaural Braincodec (512 channels) for a pure tone (0.5 kHz), an amplitude modulated tone (1 kHz carrier frequency, 4.5 Hz modulation frequency) and a speech sound coming from 3 different simulated locations (left, front 0°, right).

## Discussion

Here, we demonstrated with the Braincodec algorithm that it is possible to construct an efficient encoder of natural sounds by combining a compression and a classification neural network (**Fig. 1**). This solution transmits a high level of acoustic detail in a code largely compatible with the representations of sounds in the auditory cortex (**Figs. 2** and **3**). Several features make Braincodec an effective algorithm to concretely implement a neurostimulation implant targeting the auditory cortex. Firstly, Braincodec does not require specific training with biological data. It generates representations similar to the auditory cortex by classifying large sets of sounds into human categories, a robust phenomenon demonstrated with different types of deep networks^32,33,41,58^. An alternative approach could have been to collect a large set of human auditory cortex data and train the encoder to reproduce this data. If this approach permits good fitting performance on single cell data from animals^59,60^, it remains impractical for use in a brain implant. Training our encoding model for accurate reconstruction of any type of sounds required datasets of several thousands of hours. Despite rapid progress of single unit recordings in the human brain^61^, it is currently difficult to record thousands of hours of high resolution neural activity from thousands of human primary auditory cortex neurons. As sound reconstruction training with Braincodec shows (**Fig. 1c-d**, 30 training epochs ∼4000 hours of sound), such large datasets are necessary to guarantee robust information transmission by the encoder. Braincodec solves this issue by providing a code which can be intensively trained to its best without brain data and reproduces diverse response features observed in the auditory cortex (**Fig. 2, 3, Extended Data Fig. 3**).

The second advantage of Braincodec is that it requires a relatively low number of channels to transmit the level of acoustic information necessary for natural perception. Excellent performance is obtained with 256 to 512 channels for monaural and binaural versions of the algorithm (**Figs. 2,6**). Recent developments of high-density electrode arrays for clinical applications^62^ make these numbers easily reachable over the ∼4 cm^2^ surface area of the primary auditory cortex. Ideal stimulation arrays must permit the activation of neural ensembles that are sufficiently large to produce auditory perceptions^3^. They must also generate single electrode stimulations that are sufficiently localized to transmit independent information from each electrode. As shown in the visual and somatosensory cortex, arrays of penetrating electrodes provide an effective way to match these two constraints in human subjects^63,64^. Modern surface electrodes with small contacts could also be used as they can reach resolutions as low as 200 µm in the mouse^16,62^, which is sufficient to place up to 512 independent electrodes over a few cm^2^. However, high-resolution surface devices remain to be validated for stimulation in human trials. The use of 256 or 512 electrodes with high spatial resolution and Braincodec-based encoding guarantees that most acoustic information will be sent to the auditory cortex (**Fig. 1**). It does not permit the reproduction of precise sound driven activity patterns. Even if Braincodec’s output can be aligned to the mesoscopic organization of sound responses in the auditory cortex (**Fig. 2**), the exact microscopic organisation of sound-driven activity will not be reproduced. Therefore, the effectiveness of the cortical implant will depend on the ability of the patient’s brain to associate a new set of sound representations to meaning. We have shown in a previous study that the reformatting of the cochlear code into the cortical code for sound is necessary for efficient association of sound to behavioral responses^28^. Therefore, by efficiently mimicking this reformatting, Braincodec (**Fig. 3**) is expected to facilitate the learning process necessary for patients to use the implant. This approach is similar to the one taken with the cochlear implant, which also requires readaptation of the brain to a new input coarsely aligned to normal cochlear activity ^65^. This learning process allows many patients to reach perceptual performances close to the theoretical limits of information throughput by cochlear implant encoders, allowing speech comprehension but coarse perception of acoustic fine structure^2,65^ (see **Fig. 4 & Supplementary Videos 3-6**). Braincodec and cortical implants represent a new solution to increase the limits of the encoding stage of the implant. This paves the way for the next generation of hearing restoration solutions, which could provide perceptual performance close to natural hearing once fully developed for patients.

Lastly, Braincodec has a relatively small size of ∼4 million parameters, much smaller than deep networks for sound recognition, or even generic filter banks used as models of auditory cortex computation (**Extended Data Fig. 2**). Implementation on a low-power processor, needed for an implant, is therefore feasible. Braincodec is an algorithmically simple solution based on the combination of an autoencoder and a classifier. Autoencoders are generic tools for data compression, and classification networks were shown to produce biomimetic codes for vision and proprioception^58^. Therefore, Braincodec is likely to represent a powerful generic solution to construct encoders for sensory restoration.

## Methods

### Braincodec architecture and training

#### Network architecture

The architecture used to train the Braincodec cortical implant encoder is composed of three deep neural networks (**Fig. 1b**). (i) The encoder receives audio inputs (waveform sampled at *f_sr_* = 24 kHz) and outputs a latent representation of *N* stimulation channels (values used: 32, 64, 100, 128, 256, 512) at a framerate *F_c_* = 40 Hz. The choice of 40 Hz is based on classical estimates of the boundary between pitch and flutter in the perception of amplitude modulated sounds^20,37^ and on the common frequency limits for synchronization of cortical neurons with amplitude modulation. Output values are clipped between 0 and 50. (ii) The decoder network reconstructs the sound waveform at the original sampling frequency using as inputs the latent representation produced by the encoder. (iii) The classifier takes as input the encoder output, time-averaged over sound sample duration (2 s in our dataset) and outputs one of *M* semantic labels used to classify sound samples (*M* = *423* for our dataset). The encoder and decoder architectures are derived from the autoencoder network of Défossez et al^34^ freely available on GitHub (https://github.com/facebookresearch/encodec).

The encoder model consists of one 1D convolution layer with *C* causal kernel of size of 7 followed by *B* convolution blocks. Each convolution block is composed of a single residual unit followed by a down-sampling layer consisting in a strided convolution, with a kernel size *K* of twice the stride *S*. The residual unit contains two convolutions with kernel size 3 and a skip-connection. The number of channels is doubled whenever down-sampling occurs. A final 1-dimensional convolution layer with a kernel size of 7 and *N* output channels producing the time-dependent vector of stimulation patterns (*r_n_*(*t*))*_n_*_=_*_1_*_:*N*_ (also referred to as Braincodec’s latent space). We use *C* = *32*, *B* = *4* and (*2*,*5*,*6*,*10*) as strides. In total, the encoder is then composed of 14 1D-convolution layers. We use ELU as a non-linear activation function. We used weight normalization with statistics computed over the time dimension. The decoder model *G* mirrors the encoder, using transposed convolutions instead of strided convolutions, and with the strides in reverse order as in the decoder, outputting the final mono audio. The classifier consists of a single fully connected layer (*N*, *M*). For each sound, the predicted label corresponds to the channel with the largest output.

#### Training objective

Braincodec is trained to minimize the weighted sum *L* = *L_R_* + *δ_cl_* ⋅ *L_cl_* of the reconstruction loss *L_R_* and the classification loss with a discount factor set to *δ_cl_* = 2000.

The reconstruction loss is backpropagated through the decoder and encoder and is the sum of three terms:

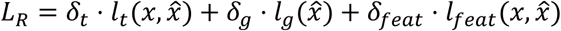

where *δ_t_* = *0*.*1*, *δ_g_* = *4* and *δ_feat_* = *4*. *x* and *x̂* are the original and reconstructed sound waveforms. To stabilize training of the autoencoder losses, we used the loss balancer introduced by Défossez and colleagues^34^ with the default parameters.

The first term minimizes the L1 distance between the original and reconstructed sound waveforms:

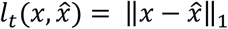

The second and third terms are adversarial loss terms aiming to minimize systematic perceptual differences between original and reconstructed sounds. They are computed thanks to 5 discriminator neural networks trained to discriminate original and reconstructed sounds based on their Short time Fourier Transforms (STFT) with 5 different window lengths ([2048, 1024, 512, 256, 128]; real and imaginary parts are concatenated). Each discriminator network is composed of a 2D convolutional layer (kernel size 3 x 8 with 32 channels), followed by 2D convolutions with increasing dilation rates in the time dimension of 1, 2, and 4, and a stride of 2 over the frequency axis. Then a final 2D convolution with kernel size 3 x 3 and stride (1, 1) is applied. The non-linear activation functions of each layer are LeakyReLU and weight normalization is applied to the network.

The 5 discriminators are trained in parallel of the encoder, decoder and classifier using the backpropagation algorithm to minimize the following hinge-loss adversarial loss function:

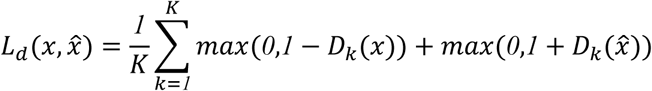

where *K*= 5 is the number of discriminators modules.

The second and third terms of the reconstruction loss are the generator loss:

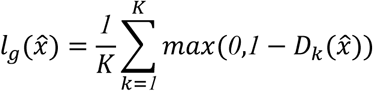

where *K* is the number of discriminator modules, and the relative feature matching loss:

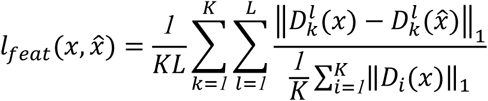

where the mean is computed over all dimensions, 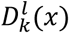 is output vector of layer *l* in discriminator module *k* for sound *x*, and *L* is the number of layers in the discriminator modules.

The classifier *φ* is trained with a binary cross entropy loss for the classifier output vector (*z_n_*)*_n_*_=_*_1_*_:*M*_ and the one-hot vector label (*y_n_*)*_n_*_=_*_1_*_:*M*_:

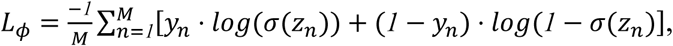

where *σ* stands for the sigmoid function. The classifier performance is evaluated as the fraction of correctly classified sounds.

#### Training

Braincodec was implemented in PyTorch^66^ and trained in two stages. In the first stage, only the encoder and decoder are trained to reconstruct sounds for 30 epochs by minimizing *L_R_*. In the second phase, reconstruction training is continued for 120 additional epochs while training the classifier to minimize *L_cl_*. Backpropagation of classification error affects both the classifier and the encoder. Backpropagation of reconstruction error affects both the decoder and the encoder. One epoch corresponds to 4,000 updates of the Adam optimizer^67^ with a batch size of 64 examples of 2 seconds each, a learning rate of *1* × *10*^−^*^4^*, and parameters *β_1_* = *0*.*9* and *β_2_* = *0*.*999*. All the models were trained using 1 Nvidia A100 or 2 Nvidia A40 GPUs. Each epoch consisted of 4,000 randomly-selected batches from the training set.

#### Datasets

To train the models, we used a modified version of the Word-Speaker-Noise dataset^56,68^. Our dataset consists of 31468 unique speech excerpts from English Spoken Wikipedia Corpora^69^ embedded in 374000 unique nonspeech background noise excerpts from Audioset strong^70^. Randomly sampled pairs of 2 s speech and noise excerpts were combined to yield a training dataset of 5796000 examples. A validation set of 4000 examples was similarly constructed from speech and noise excerpts excluded from training. Each example was labeled with the talker that produced the speech utterance and the word that appeared in the middle of the utterance (i.e., that overlapped the 1 s mark of the 2 s utterance). The resulting dataset contains 310 unique talker labels, 263 unique word labels and 423 unique semantic labels (such as mechanisms, wind, conversation, laughter). The main difference with the original Word-Speaker-Noise dataset, is the use of AudioSet strong instead of AudioSet^71^. The classifier was trained only to recognize semantic labels.

To evaluate reconstruction performance on an unrelated dataset in **Fig. 4c** and the decorrelation of sound representations in **Fig. 3c**, we used the ESC-50 dataset, a labeled collection of 2000 environmental audio recordings suitable for benchmarking methods^72^. The dataset consists of 5-second-long recordings organized into 50 semantic classes (with 40 examples per class) loosely arranged into 5 major categories: Animals, natural soundscapes & water sounds, human and non-speech sounds, interior/domestic sounds and exterior/urban noises.

### Braincodec variants

#### Tonotopically organized variant for the mouse implant prototype

The 100-channels model used for discrimination tests in mice produces stimulation patterns organized on a 2-dimensional grid (10 x 10) spatially structured along a tonotopic axis aligned to one of the grid diagonals (**Fig. 5**) thanks to a supplementary loss term 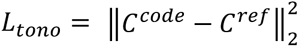 in which *C^code^* is the center-of-mass of the 2-dimensional stimulation pattern (encoder output mapped to the 2D-grid) and *C^ref^* is the location on the tonotopic axis of the peak frequency *f* in the FFT of the sound. Using the 8 log-spaced anchor frequencies 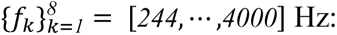

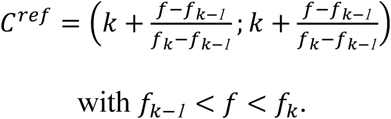

The total training objective *L* = *L_R_* + *δ_cl_* ⋅ *L_cl_* + *δ_tono_* ⋅ *L_tono_* with *δ_tono_* = 10, was minimized after the 20 training epochs dedicated to sound reconstruction alone. The Braincodec architecture used to generate tonotopic stimulation patterns included two additional recurrent neural network layers (LSTM) before the last convolutional layer.

#### Binaural Braincodec

To process spatialized binaural inputs [*x_L_*(*t*); *x_R_*(*t*)], the first layer of the encoder and last layer of the decoder are constructed with 32 kernels of size 2×7 summing the 1D-convolutions with *x_L_*(*t*) and *x_R_*(*t*). The rest of the decoder and encoder architectures are unchanged. Sounds are numerically spatialized in an anechoic environment using the Virtual Acoustic Room simulator of Saddler and colleagues^56^. Given the source angular position (θ,*φ*), [*x_L_*(*t*); *x_R_*(*t*)] were obtained by convolving the sound waveform *x*(*t*) with the Head-Related Impulse Responses of the left and right ears (*ℎ_L_*(*θ*, *φ*) and *ℎ_R_*(*θ*, *φ*)). The Source angular positions were defined conventionally from the antero-posterior axis of the head (0° for sources oriented to the face) on a grid ranging from −90° to 90° with 5° intervals for azimuth (θ) and from −40° to 60° with 10° intervals for elevation (ϕ). During training, we randomly assigned source direction on the grid to the 5796000 sound samples of the Braincodec training set to construct the main training set (70% of the total training set). To ensure that the model effectively decouples the left and right channels 20% of the training samples were monaural sounds (one of the two channels selected at random). To avoid lateralization biases, 10% of the training samples were a selection of samples from the main training set for which the left and right channels are swapped.

The classifier network is extended with a fully connected linear layer mapping the time-averaged output vector of the encoder to a 3-dimensional vector *v* corresponding to the predicted 3-dimensional location of the source. The prediction is then restricted to the sound source direction by normalizing *v* with its L2-norm and training loss for sound direction is:

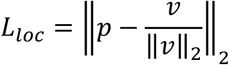

where *p* represents the position of the simulated sound source on the unit sphere.

Overall, the binaural braincodec training objective was

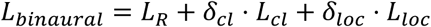

where *δ_loc_* = *100* and *δ_clf_* = *10000*. The training of binaural Braincodec happened in two stages with 40 epochs dedicated to sound reconstruction only (*L_R_*), followed by 120 epochs during which all training objectives are optimized.

### SI-SNR

We assessed the similarity of the original *x* and reconstructed *x̂* waveforms with the scale-invariant signal-to-noise ratio (SI-SNR)^42^. The sounds are centered around their mean (*x̂_c_*, *x_c_*), and the original sound is scaled to the reconstruction:

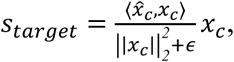

where ⟨⋅,⋅⟩ denotes the inner product over the time axis, ‖. ‖_2_ the Euclidean norm, and *ε* a small constant (scaled with the sound length) added for numerical stability.

The SI-SNR is defined as:

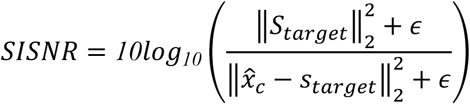

### Comparison with human brain data

We compared Braincodec stimulation patterns with a dataset of auditory cortex responses to 165 natural sounds measured with blood-oxygen-level-dependant functional magnetic resonance imaging (BOLD fMRI) by Norman-Haigniere et al.^43^, and freely available at https://mcdermottlab.mit.edu//tuckute_feather_2023/data.tar. The comparison methodology is adapted from Kell et al.^32^ and Tuckute et al.^33^. Summary similarity measures were based only on voxels of the primary auditory regions (areas Te 1.0, Te 1.1 and Te 1.2 defined in the Julich Brain Atlas v3.1^73^ aligned on FreeSurfer FsAverage-7). We first identified for each voxel *j* the linear transformation *w_j_* that best fits voxel responses based on the *N* channel encoder outputs for *s* = 83 training sounds using L2-regularized (“ridge”) regression:

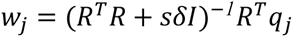

where *w_j_* is a *N*-length vector, *q_j_* is an *s*-length vector containing the voxel’s mean response to each sound, *R* is the *N* × *s* matrix of time-averaged encoder outputs and *I* is the *N* × *N* identity matrix. The prediction of sound responses is then computed as *q̂_j_* = *Rw_jj_*. We used 83 sounds to train the regression and representation similarity is measured on the 82 remaining sounds, taking the median across 10 training/test cross-validation splits. To determine the regularization coefficient *δ* for each training split, we performed the regression on 82 of the 83 training sounds for 100 logarithmically spaced values [1e-50, 1e-49,…, 1e48, 1e49] of *δ* and selected the value minimizing the mean square error on the remaining sound.

We then evaluate the similarity between the actual sound responses and their model-based prediction with a noise-corrected measure of variance explained:

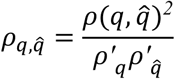

where *q* is a voxel response to the 82 left-out sounds averaged over the 3 recorded scans, *q̂* = *Rw* is the predicted response to the 82 left-out sounds (with regression weights *w* learned from the other 83 sounds), *ρ* is the Pearson correlation coefficient, *ρ′_q_* is the estimated reliability of that voxel’s response to the 83 sounds, and 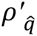 is the estimated reliability of that predicted voxel’s response. *ρ′_q_* is the median of the correlation between all 3 pairs of scans, which is then Spearman–Brown corrected to account for the increased reliability that would be expected from tripling the amount of data^74^. Note that for very noisy voxels, this division by the estimated reliability can be unstable and can cause for corrected variance explained measures to exceed one. To reduce the impact of this problem, we limited both the reliability of the prediction and the reliability of the voxel response to be greater than some value *k*^75^. For *ρ′* and *ρ′* we used *k*’s of 0.182 and 0.183 respectively^33^.

### Spatial mapping of Braincodec patterns

To illustrate how Braincodec patterns can be mapped to the primary auditory cortex (**Fig. 2e**) based on functional maps, we used the *N* x *Q* matrix 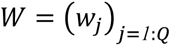 which gathers the linear predictors of voxels activity averaged across 10 cross-validation splits, with *Q* the number of voxels on the 2-D cortical surface of a target rectangular area (size *22* × *37* for the right hemisphere, *24* × *20* for the left hemisphere) covering a primary auditory cortex. This allows us to compute *Q* stimulation time courses *r̂*(*t*) = *Wr*(*t*), spatially mapped to the recorded voxels, based on the *N* stimulation time courses *r*(*t*) output by the Braincodec encoder. *r̂*(*t*) is then downsampled to *N* = *512* channels (*16* × *32* electrode array) by linear interpolation.

### Cochlear implant encoder simulation and evaluation

Simulation of the cochlear implant encoders was performed with the Nucleus Toolbox in MATLAB^76^, using the advanced combination encoder (ACE) algorithm^50^ for a 22-channel electrode array with sounds sampled at 24 kHz. ACE was parameterized with conventional values including an output frequency of 1 kHz, a sliding window length for the Fast Fourier Transform (FFT) of 8 ms (128 samples), a sliding window step of 1 ms, and 12 out of 22 channels selected at each step based on the FFT-power *E_u_* in the frequency band *u* associated to the channel (only the 12 channels with the highest power have non-zero output on each time step). The output level for each channel is obtained after log-compression of the FFT power 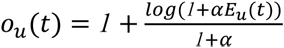 where *α* > *0* is the loudness-growth-function scaling factor (the LGF-alpha) is usually derived from the patient’s clinical map, but here with the default clinical parameters 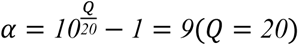.

To evaluate information loss in ACE, we used a deep-neural-network synthesis building directly on the reconstruction pipeline introduced in the “Braincodec neural network architecture and training” section. Instead of feeding the model raw waveforms, we use the output *o_u_*(*t*) or intermediate *E_u_*(*t*) (before channel selection) representation generated by ACE as inputs of a decoder network (one separately trained decoder for *o_u_*(*t*) and for *E_u_*(*t*)). The decoder is parameterised to match the dimensions of the cochlear-implant codes. The first layer learns 512 filters of dimension 7×22. It is followed by four blocks of three convolutional layers with strides (*3*,*2*,*2*,*2*) and a last layer, which progressively up-sample the input feature map back to the original audio sampling grid. During training the output of the decoder is compared with the corresponding ground-truth waveform using the objective function *L_R_* described above.

Deep neural network synthesis was compared to a vocoder which construct a sound based on the FFT power values *P̂_band_* (*t*,*m*) of the 22 frequency bands according to the formula:

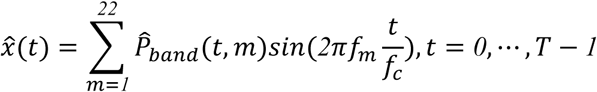

with *F* the number of analysis frames, *f_c_*=16 kHz the audio-output sampling frequency and *f_m_* the centre frequency of band *m*.

### Behavioural experiments

#### Animals and housing

Every experiment involving animals has been carried out following a protocol approved by an ethical commission, in accordance with the French Ethical Committee (Direction Générale de la Recherche et de l’Innovation) and European legislation (2010/63/EU). Procedures were approved by the French Ministry of Education in Research after consultation with the ethical committee #59 (authorisation number APA-FIS#28757-2020122111048873). Optogenetic experiments were conducted on adult Emx1-IRES-Cre x Ai27D (Digital Micromirror Device, head-fixed experiments) and C57Bl6J or CamKII-CreERT2 x Ai228 (µLED array, freely moving experiments) mice group-housed in an animal facility (08:00-20:00 light) at a temperature of 22 ± 2∘C and humidity between 60–65%, in cages with access to enrichment (running wheel, cotton bedding and wooden logs).

#### Cranial window or μLED array implantation surgery

Procedures were performed in 8–12-week-old mice. Buprenorphine (Vertergesic, 0.05–0.1 mg/kg) was given 30 min pre-anaesthesia; isoflurane was induced at 3% and maintained at 1–1.5% on a heating blanket maintaining the mouse at 37°C. Eyes were protected with Ocry gel (TVM Lab) and xylocaine (20 mg/ml; Aspen Pharma) was injected at the incision site. The skin was removed above the dorsal part of the skull and a layer of SuperBond (Sun Medical, Shiga, Japan) was deposited on the cleaned skull surface. The right masseter was partially removed. For µLED array implantation a 2.4 × 2.4 mm rectangular craniotomy was made over the right auditory cortex using skull sutures as landmarks. In two C57Bl6J mice, AAV vectors encoding ChRmine under the control of the CaMKIIα promoter (pAAV-CamKIIα-ChRmine-mScarlet-Kv2.1-WPRE (AAV8). Addgene number: 130991) were injected (6 to 8 injections of 150 nL) to drive expression in pyramidal neurons of the auditory cortex in two mice. In three CamKII-CreERT2 x Ai228, ChRmine was constitutively expressed after three intraperitoneal Tamoxifen injections at 6 weeks, and no virus injection was performed. Then, the μLED array was placed on the cortical surface, glued to the skull (SuperGlue, Loctite), and the flexible PCB secured with dental cement (Ortho-Jet, Lang). For cranial window implantation, a ∼5mm diameter craniotomy was performed and a 5mm glass coverslip was positioned and sealed with Superbond. For reward delivery in the head-fixed DMD experiment, a bipolar stimulation electrode (60 µm diameter, twisted stainless steel; PlasticsOne) was implanted in the medial forebrain bundle (MFB) using stereotaxic coordinates (AP −1.4, ML −1.2, DV +4.8). The electrode and the head plate were then fixed to the skull using the Ortho-Jet dental cement. For all experiment experiments, a head bar was fixed with Ortho-Jet dental cement. Post-op, mice received s.c. glucose (G30) and meloxicam (Metacam, 1 mg/kg), then recovered for 1 week with Metacam in drinking water or DietGel (ClearH2O). Mice were group-housed; rare temporary isolation occurred if cage-mates chewed device connectors.

#### Sound generation and delivery

Sounds were generated with Matlab (The Mathworks, Natick, MA) and were delivered at 192 kHz with a NI-PCI-6221 card (National Instruments) feeding an amplified free-field loudspeaker (SA1 and MF1-S, Tucker-Davis Technologies, Alachua, FL) positioned 15 to 20 cm from the mouse ear. Sound intensity was cosine-ramped over 10 ms at the onset and offset to avoid spectral splatter. Experimental setups were isolated from external noise sources by sound-proof boxes (Decibel France, Miribel, France) providing 30 dB attenuation above 1 kHz. Sounds were calibrated in intensity at the location of the mouse ear using a probe microphone (Bruel & Kjaer, type 4939-L-002). For pure tone discrimination experiments, sixteen 70dB SPL pure tones were generated with frequencies logarithmically spaced from 4 to 16kHz.

#### Braincodec pattern generation and delivery

We designed and fabricated 100-pixel arrays of blue μLEDs (450 nm), each integrated to a flexible PCB (total mass <300 mg) for chronic implantation and optogenetic stimulation of the mouse auditory cortex^18^. Note that 450nm light efficiently activates ChRmine because of its broad sensitivity. The 10 x 10 array of μLEDs is matrix-addressed via 10 common anodes and cathodes, minimizing interconnection. Devices were encapsulated with a 15 μm layer of parylene-C ensuring device stability over weeks-long chronic experiments. A miniaturized connector and lightweight cable (Omnetics Connector corp.) permitted interconnection from the μLED array to a driver system while allowing the mouse to behave freely. The driver system incorporates a PYNQ-Z2 board (AMD) and custom shield PCB housing the μLED driver circuitry (**Extended Data Fig. 5**). This includes a 10-channel current DAC (Analog Devices LTC2662) connecting to the μLED array anodes, and a 10-channel solid-state switch array (TI TMUX7211) allowing the common cathodes to be connected to ground. Using time-division multiplexing, this design allows arbitrary pattern generation, albeit at a 10% duty cycle. Drive current is automatically inferred from the requested cortical irradiance using optical measurements made before device implantation. This allows the current to be individually calibrated for each μLED, accounting for small variations in the fabrication process. The μLED array driver firmware runs on a Microblaze processor (AMD) in the programmable logic of the Zynq 7020 SoC on the PYNQ-Z2 board. The system receives a TTL trigger input for experimental synchronization. A Python interface (JupyterLab) allows the user to operate the system for straightforward loading of Braincodec patterns.

We generated 16 Braincodec spatiotemporal patterns corresponding to pure tones logarithmically spaced from 500 to 2000 Hz (**Fig. 5a**). We used the tonotopically organized variant of Braincodec for a 10×10 rectangular array, trained on natural sounds whose frequency spectrum rarely extends above 4 or 8 kHz. Therefore, Braincodec cannot properly encode sounds in the mouse hearing range (∼ 4 - 80 kHz). Our experiment therefore tests the perception of patterns generated to cover 2 octaves in the human frequency range to be compared to perceptual discrimination of real pure tones covering 2 octaves in the mouse frequency range. Braincodec stimulation patterns amplitudes were mapped to the peak µLED irradiance interval [0, 10 mW/mm^2^] to match the maximum output of the µLED array. The µLED array was placed on the auditory cortex such that the tonotopic axis of the stimulation patterns matches roughly the axis of the main tonotopic field A1.

#### Tonotopic patterns generation and delivery

To compare Braincodec stimulations to a simpler alternative model, we measured perception of 8 optogenetic stimulations corresponding to rectangular stripes (560 x 125 µm) whose long axis was perpendicular to the main tonotopic axis of A1 and which were regularly spaced from the ∼low to the high frequency end of the axis (**Extended Data Fig. 7c,d**). The tonotopic axis was identified with intrinsic imaging under light isoflurane anaesthesia (1%; SomnoSuite, Kent Scientific) on a thermal blanket. Imaging was performed through a cranial window implanted 1–2 weeks earlier, using a CCD camera (GC651MP, Smartek Vision; 4-pixel binning; field of view 3.7 × 2.8 mm; 164 × 124 binned pixels; 5.58 µm/pixel; 20 fps) operated by a custom MATLAB software. Reflectance signals were recorded under 780 nm LED illumination (Thorlabs), and vasculature images were acquired under 480 nm illumination (Conrad). Two-second sequences of pure tones (80 dB SPL; 4, 8, 16, 32 kHz) and white noise were presented every 30 s (10 trials per sound). For each sound, baseline (3 s pre-onset) and response (3 s post-onset) images were computed, and δR/R0 was averaged across repetitions and smoothed with a 2D Gaussian filter (σ = 4 pixels). The posterior response peak for 4 kHz and dorsal response peak to 16 kHz usually defined the low and high frequency ends of A1’s tonotopic axis as responses to 32 kHz were often weak and are in fact located at a similar position as responses to 16 kHz (**Extended Data Fig. 7d-f**). Axis length ranged from 1 to 1.2 mm and was divided into 8 equally spaced locations at which the stimulation stripes were positioned. We observed that rectangular stripes finer than 125 µm are poorly perceived by mice and therefore restricted the number of stimuli for catch trial experiments to 8 non-overlapping stripes over ∼1 mm (2 target and 6 catch trial stimuli). Go and NoGo stimuli were assigned to the 4-kHz and 16-kHz ends, respectively. Patterns were exported as BMPs and converted to MP4 videos for playback using custom MATLAB code, and were repositioned at the same location across days using the surface blood vessel pattern, through a custom alignment routine written in Matlab.

Light patterns (0.5 s duration with 25 ms on and off illumination periods at 20Hz) were delivered by a digital micromirror device (DLP LightCrafter, Texas Instruments) illuminated by a 460 nm LED. To project a two-dimensional image onto the auditory cortex surface, the image of the micromirror chip was collimated through a 150 mm cylindrical lens (Thorlabs; 2-inch diameter) and focused through a 50 mm objective (NIKKOR, Nikon). Light collected by the objective passed through a long-pass dichroic beam splitter (>640 nm; FF640-FDi01, Semrock) and was captured by a CCD camera (GC651MP, Smartek Vision) equipped with a 50 mm objective (Fujinon HF50HA-1B, Fujifilm).

#### Behavioral training procedures

Sound and tonotopic pattern discrimination tasks were performed in head-fixed mice with water rewards (solenoid-controlled 4 µL drops) for sounds and MFB stimulations (0.5V to 3V; biphasic pulses of 1ms to to 10ms for 100ms depending on mouse’s response) for tonotopic patterns. It was previously shown that these two reward protocols yield identical auditory psychometric measures^77^. Braincodec pattern discrimination and associated sound discrimination tasks were performed in a freely moving setting with water rewards (4 µL drops). Mice rewarded with water were under a water restriction protocol (1 mL per day during weekdays and access to 4% citric acid in water during weekends). MFB rewarded mice received food and water ad libitum. In the head fixed setting, the head post was attached to a metal bar and the mouse was resting in a crouching position in a custom 3D-printed plastic tube with a conductive metal floor, facing a metal licking spout. Tongue contacts on the licking spout closed a 5V-powered electrical circuit between the spout and the floor producing voltage deflections across a series resistor measured to count individual licks^78^. Freely moving experiments were carried out in a transparent polypropylene box (27 x 25 x 22 cm) with an infrared light barrier in front of the water delivery spout which detected when the animal’s head was positioned for licking. Beam break duration was used as a proxy for licking.

In all tasks, mice performed a single 1 to 3 hours training session per day (300-400 trials with water reward, 300–1000 trials with MFB reward). A trial consisted of uniformly distributed random inter-trial intervals (1-2s for water reward, 6-8s for MFB reward) and ‘no lick’ period (1-2s for water reward and 3-5s for MFB reward). During the latter, any lick delayed the start of the trial by a supplementary ‘no lick’ period. After the ‘no-lick’ period, the 0.5 s stimulus was played. The response window (1s for water rewards, 1.5s for MFB rewards), during which licking was measured, started with the stimulus onset. Reward was delivered in Go trials if licking during the response window was above a threshold (5 licks or beam beak for 75% of the response window duration for water rewards; 1 lick for MFB reward). In NoGo trials, a random timeout (3-4 s for water rewards, 5 - 7 s for MFB rewards) was inserted after the response window if licking was above the same threshold.

For all tasks, the first session aimed at habituating mice to lick on the waterspout, by rewarding them immediately after licking on the spout or breaking the infrared beam. In the second session, mice started a Go-stimulus association training, in which the Go stimulus was presented in 80% of the trials while the remaining trials were blank stimuli with no reward to probe baseline licking behavior. When mice reached 80% correct Go trial responses during a session, the NoGo stimulus was introduced in the next session to start the actual stimulus discrimination training. For the task with MFB reward, the blank stimulus was maintained during discrimination training and for all tasks with water rewards, it was removed. Once stimulus discrimination was acquired (at least one session with 80% correct performance), non-rewarded catch trials occurring with a 20% probability were randomly interleaved within Go and NoGo trials to measure sound categorization accuracy.

The Go-stimulus association and discrimination training phases for optogenetic stimulation were started with more salient stimuli for faster training^79^. For the Braincodec pattern discrimination, the first Go and NoGo stimulation consisted of 3 × 3 μLED patterns (20 Hz, 50% duty cycle, 10 pulses) positioned at opposite corners of the 10 × 10 matrix. The irradiance at the cortical surface was set to 10 mW/mm^2^. Once the discrimination task was acquired the irradiance for these patterns was progressively reduced to 1 mW/mm^2^. Discrimination training with Braincodec was started after mice showed more than 80% correct performance at this low irradiance for the square patterns. For the tonotopic patterns, the association and discrimination phases started with 400 x 400 μm squares gradually reduced to 560 x 125 µm stripes when mice reached 80% correct performance.

In all optogenetic tasks, a white LED lamp was directed at the mouse’s eyes to reduce visual perception of optogenetic stimuli. Also, for all tasks, at the beginning of each session, ∼10 trials with ‘free rewards’ were given independently of the licking to motivate the mouse. These trials were not included in quantifications.

## Data availability

All data acquired in this work are available from the corresponding authors upon request.

## Code availability

All codes developed in this work are available from the corresponding authors upon request.

## Acknowledgements

We acknowledge the support of the HPC Core Facility of the Institut Pasteur, and of the Animal Facility and Signal Processing and Acquisition Facility for this work. We thank Kathleen Cho, Sophie Bagur, Tania Rinaldi Barkat, James Taylor, Manvi Agarwal, Etienne Thoret and Pascal Belin for their feedback on the manuscript. We thank Matteo Pisoni for his help with mouse preparation. We also thank Aymard Grou-Radenez, Clément Lefebvre and Alicia Aniorte García for their early contributions.

## Author information

These authors contributed equally: Charly Lamothe, Antonin Verdier, Corentin Caris.

These authors jointly supervised this work: Brice Bathellier, Romain Serizel and Keith Mathieson.

## Contributions

Conceptualization: C.L., A.V., K.M., R.S. and B.B. Methodology—algorithm: C.L., A.V., C.C., R.O., R.S.R. and B.B. Methodology—implant R.G., E.B. and K.M. Methodology—behaviour A.V. and B.B. Investigation—algorithm: C.L., A.V., C.C. and R.O. Investigation—behaviour: A.V., E.M.K., P.P., R.G., E.C., S.C. and B.B. Formal analysis: C.L., A.V., C.C., R.O. and B.B. Resources: R.G., R.S.R, S.C., E.B., K.M., R.S. and B.B. Software: C.L., A.V., C.C. and R.O. Visualization: C.L., A.V., R.G. and B.B. Supervision: C.L., A.V., K.M., R.S. and B.B. Writing—original draft: C.L., A.V., R.G., R.S. and B.B.

## Extended data figures and tables

**Extended Data Fig. 1:**
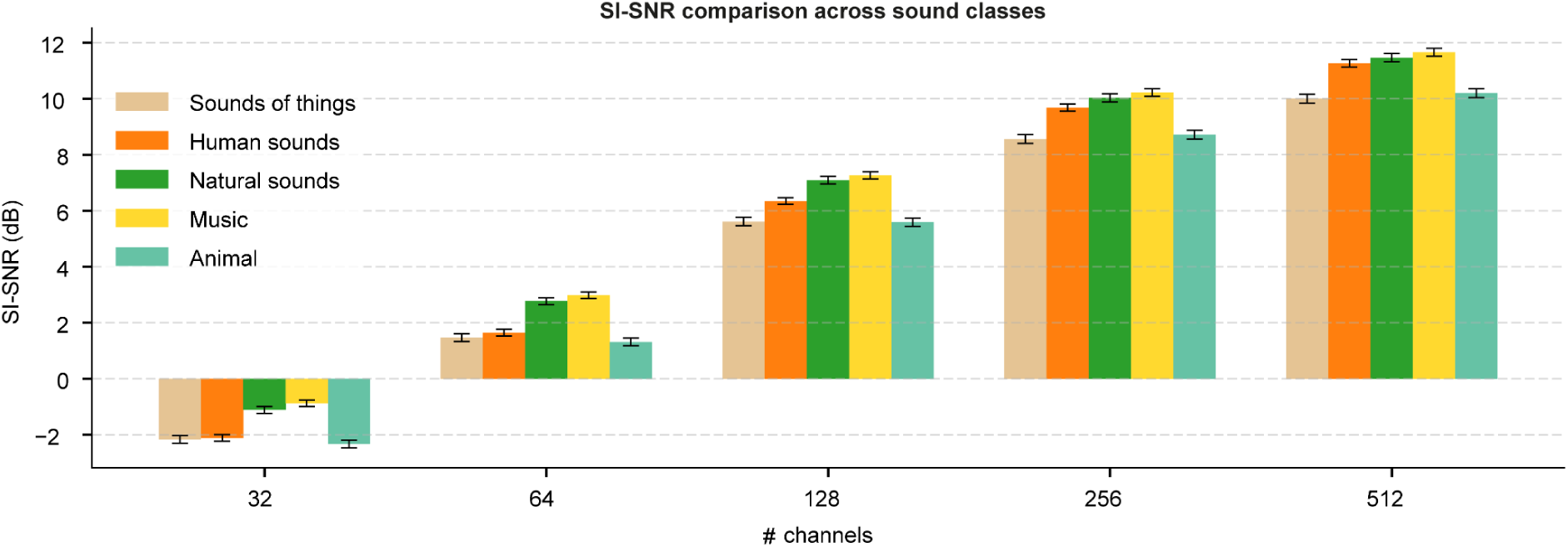
Category-specific reconstruction fidelity of Braincodec models. SI-SNR (dB) between original 24 kHz waveforms and their Braincodec reconstructions is shown for five broad AudioSet-Strong parent categories (colours). For each category, 2,000 sound excerpts were randomly selected and their leaf-level AudioSet-Strong labels were mapped to their root ontology class. The x-axis indicates the number of latent channels in each Braincodec model (32, 64, 128, 256, 512). Bars represent mean SI-SNR and error bars denote ±1 standard error of the mean (s.e.m.) across the 2,000 excerpts per category. Higher SI-SNR values indicate better reconstruction quality. SI-SNR values are reported as the negative of the SISNR loss, corresponding to conventional SI-SNR in dB.

**Extended Data Fig. 2:**
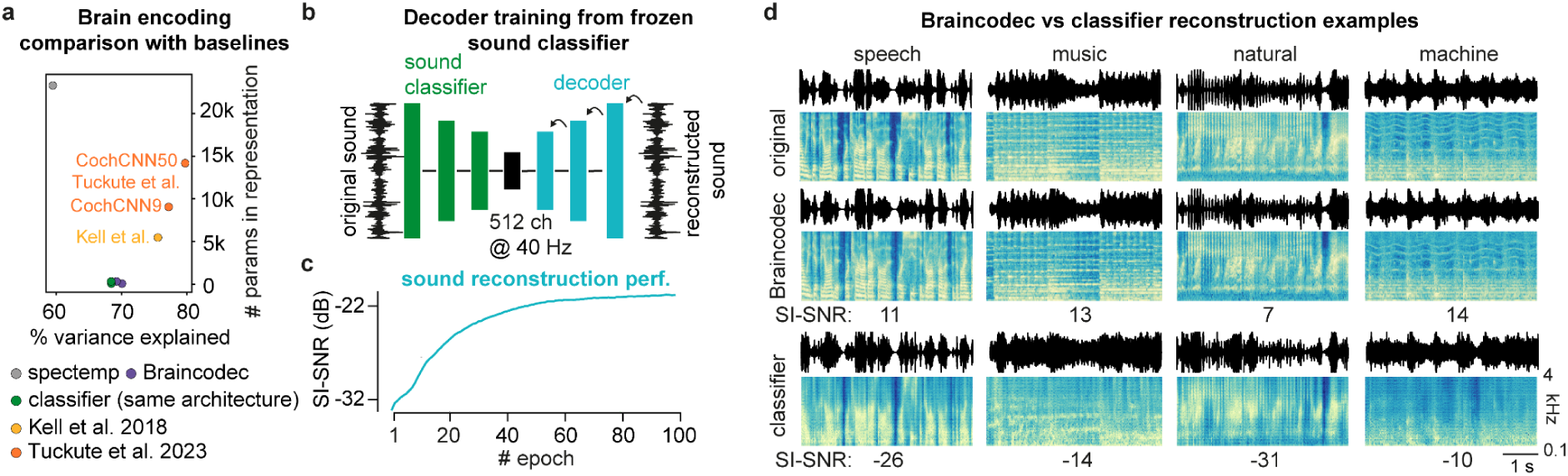
Comparison of Braincodec with baselines. **a.** Fraction of explained variance as a function of the number of parameters in each learned representation for Braincodec, a classifier (using the same backbone architecture as the Braincodec encoder), the spectrotemporal-filter model, Kell et al.’s classifier^32^, and Tuckute et al.’s classifiers^33^. **b.** Training a deep-network decoder to reconstruct sounds from a frozen sound classifier. First, the sound classifier is trained to predict semantic labels. Next, the representations of the 5.8M sounds are extracted from the penultimate layer of the classifier. Finally, the decoder is trained to reconstruct the sounds by minimizing the difference between the original and reconstructed waveforms. **c.** Mean SI-SNR of reconstructed sounds (validation dataset). **d.** Spectrogram of example sounds and their reconstructions obtained with 512 ch Braincodec and with the frozen classifier decoding.

**Extended Data Fig. 3:**
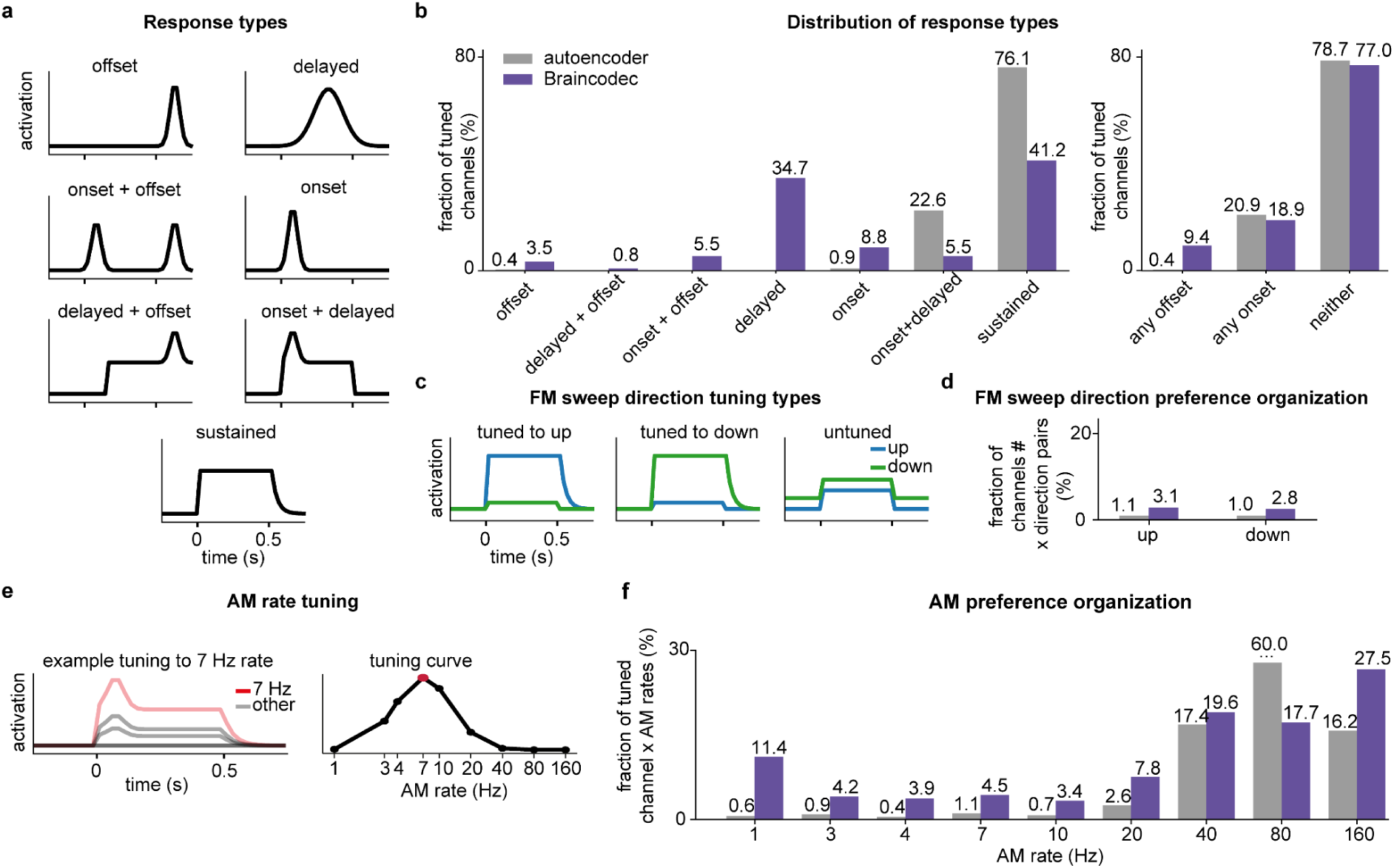
Temporal response dynamics of Braincodec. **a.** Idealized single-channel response profiles illustrating the classification scheme used to categorize pure-tone–evoked responses. **b.** Channels from the autoencoder and Braincodec models were classified based on their activity traces averaged across 128 pure tones (32 frequencies logarithmically spaced 100 Hz to 2,000 Hz, each presented at 60, 70, 80, and 90 dB SPL). Response strength was quantified in three windows relative to stimulus onset: onset (0–0.15 s), delayed (0.15–0.5 s), and offset (0.5–0.75 s). A channel was assigned to an exclusive category (delayed, onset, offset, onset+offset, onset+delayed, offset+delayed, sustained, or silent) by determining which windows exceeded the next strongest by ≥1.5-fold and passed an absolute activity threshold of 0.3. (left) Proportion of responsive channels (silent channels excluded) in each exclusive temporal category. (right) Proportion of all channels in each model showing an onset-related response (onset, onset+offset, or onset+delayed), an offset-related response (offset, onset+offset, or offset+delayed), or neither. Chi-square tests revealed significant differences between autoencoder and Braincodec models across temporal response categories (left: χ² = 327.1, p = 1.25 × 10⁻⁶⁷, n = 946 responsive channels; right: χ² = 42.3, p = 6.70 × 10⁻¹⁰, n = 1024 channels (512 per model)). **c.** Idealized single-channel response profiles illustrating the classification scheme used to categorize frequency-modulated (FM) sweep–evoked activity. **d.** Direction selectivity for frequency-modulated (FM) sweeps. FM stimuli consisted of all 120 unique frequency pairs drawn from 16 logarithmically spaced frequencies between 100 and 2000 Hz, with each sweep presented at 60, 70, 80, and 90 dB SPL (sweep duration: 0.5 s). Direction selectivity was classified using a ratio threshold of 1.5-fold between upward and downward sweep responses, with a silence threshold of 3% of the 99th percentile response magnitude. Bars show the fraction of responsive neuron × direction pairs classified as upward-(up) or downward-selective (down), aggregated across all frequency pairs and intensities. Silent channel–direction pairs are excluded from the quantification. The numbers above each bar indicate the fraction of responsive pairs. Braincodec showed a significantly higher proportion of direction-selective neurons compared to the autoencoder (χ² = 1026.3, p = 3.37 × 10⁻²²⁵, n = 108,893 responsive neuron × direction pairs). **e.** (left) Idealized response profile example of a channel tuned to AM rate. (right) Corresponding AM tuning curve. **f.** Distribution of AM rate tuning for the autoencoder alone and for Braincodec. Stimuli were amplitude-modulated pure tones with carrier frequencies logarithmically spaced between 100 and 2000 Hz (32 frequencies), presented at 60, 70, 80, and 90 dB SPL, and modulated at 1, 3, 4, 7, 10, 20, 40, 80, or 160 Hz. Channel × condition pairs were classified as silent (maximum response < 3% of 99th percentile), flat/untuned (all AM rates within 0.5–1.5 × mean response), or tuned (peak response at a specific AM rate). Bars show the fraction of responsive, AM-tuned channel & carrier frequency pairs for which the peak response occurred at one of the selected AM rates. Silent and untuned (flat) neuron–condition pairs are excluded. Numbers above the bars indicate the fraction of all tuned pairs. Braincodec showed a significantly higher proportion of AM-tuned responses compared to the autoencoder (69.2% vs 10.7%, p < 10⁻³⁰⁰), and the distribution of preferred AM rates differed significantly between models (χ² = 5677.4, p < 10⁻³⁰⁰, n = 45,841 tuned neuron × condition pairs).

**Extended Data Fig. 4:**
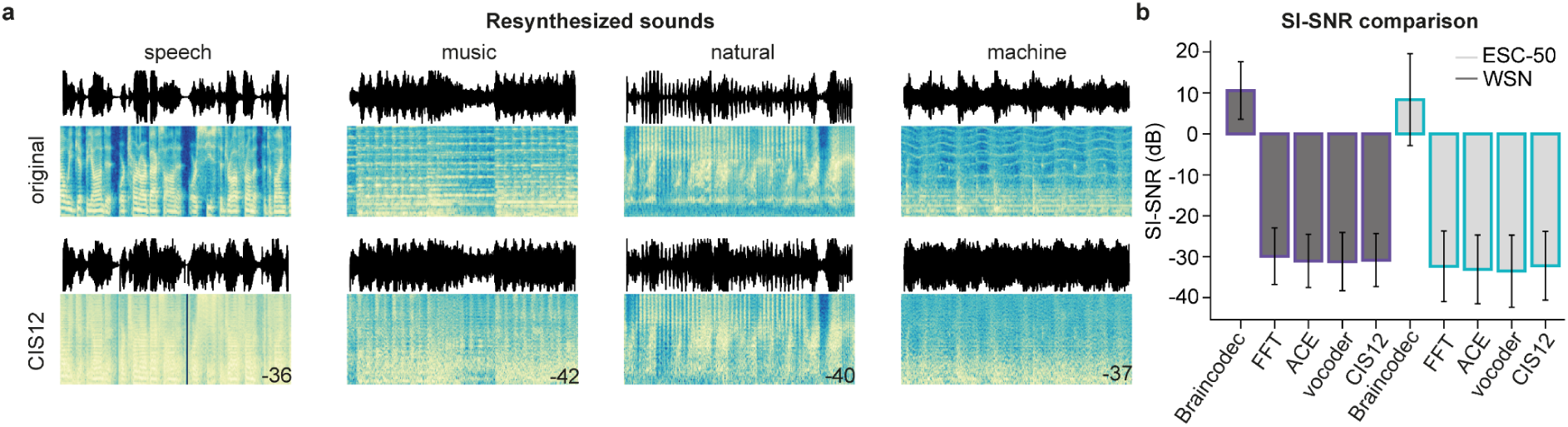
Braincodec outperforms CIS cochlear implant encoding with 12 channels. **a.** Spectrograms and sound waves for different sounds and their reconstruction by CIS12 cochlear implant reconstruction strategy. The number in the bottom right corner of the reconstructed spectrogram is the SI-SNR value indicating reconstruction accuracy (10 excellent, 0 to −10 poor, <-10 very poor, see Fig. 1c). **b.** Average SI-SNR of reconstructed sounds for the evaluation (WSR) and validation datasets (ESC-50).

**Extended Data Fig. 5:**
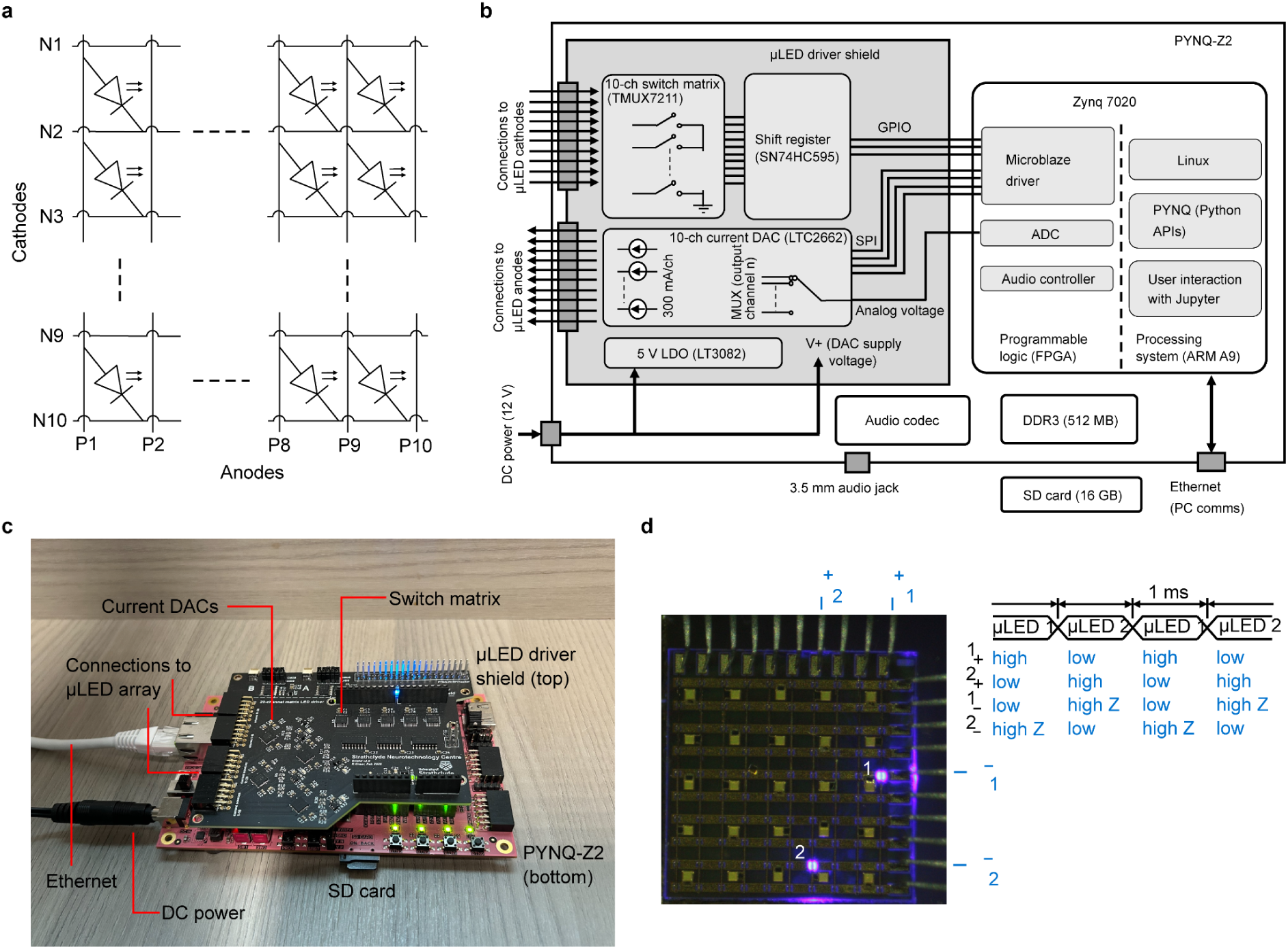
Electronic controller for the implantable µLED array. **a.** Electrical schematic showing matrix-addressed layout of µLEDs. For clarity, the schematics and labelling for µLED pixels correspond to the “top-down” view, i.e., the light emission is “into” the page. **b.** High-level schematic of electronic control system based on a PYNQ-Z2 development board and custom current driver shield PCB. The system is programmed in C (for the low-level Microblaze driver) and Python using the PYNQ framework. The Microblaze driver cycles through rows of the matrix, closing a single switch to ground and activating each DAC at the required current level. Each row is active for 10 ms with a duty cycle of 10% for the entire matrix. The current level is set in DDR3 memory by the user in a Jupyter notebook and accessed by the Microblaze through direct memory access (DMA). This allows arbitrary stimulation patterns and brightness, giving the flexibility to drive Braincodec patterns. **c.** Annotated photograph of electronic control system. **d.** Diagram showing time-division multiplexing used to achieve arbitrary control of patterns and µLED brightness.

**Extended Data Fig. 6:**
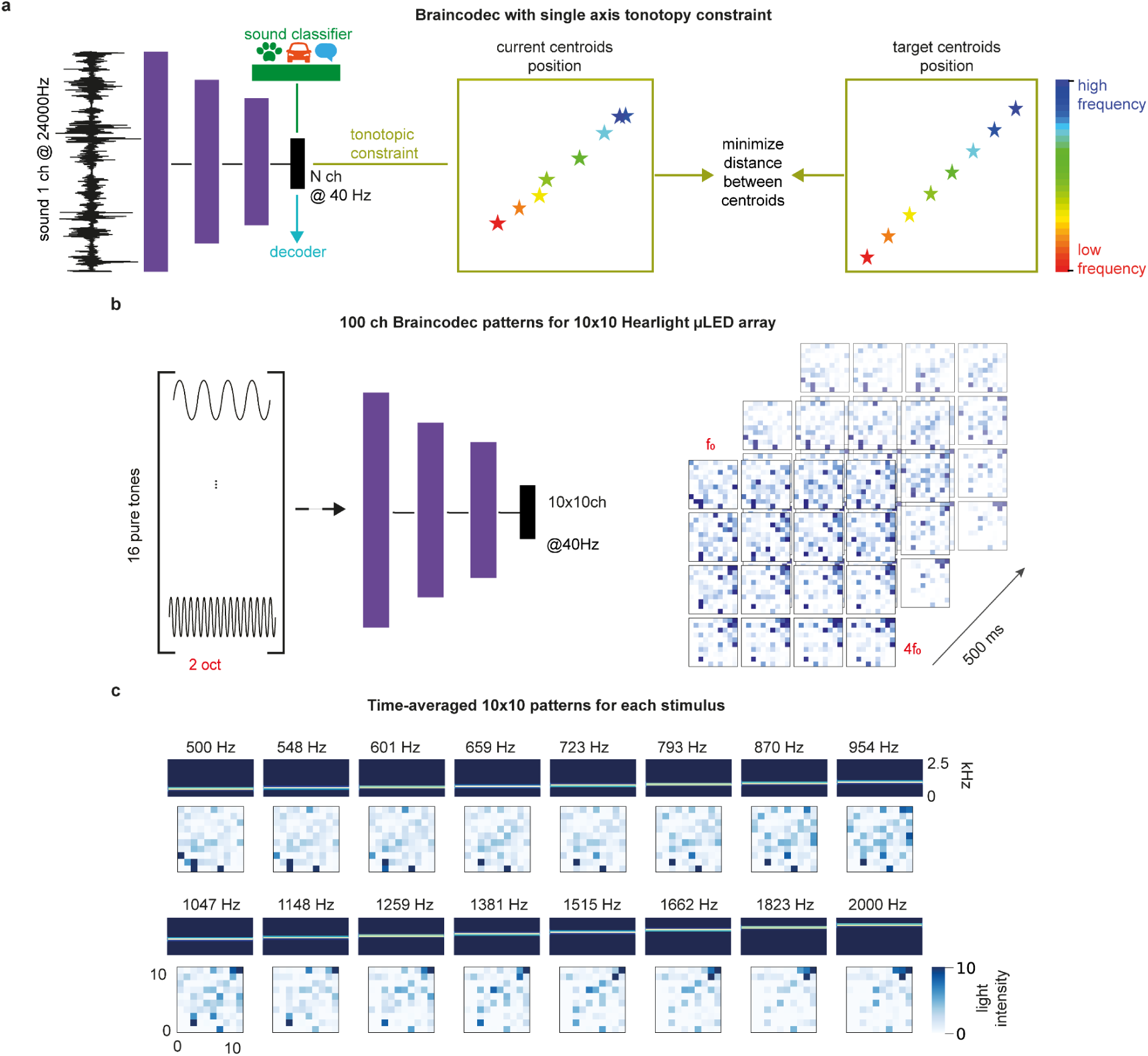
Tonotopically-organized Braincodec variant for the mouse implant prototype. **a**. The 100-channels model used for discrimination tests in mice produces stimulation patterns organized on a 2-dimensional grid (10 x x 10),) spatially structured along a tonotopic axis aligned to one of the grid diagonals (Fig. 5) thanks to a supplementary loss term that minimizes the distance between current sound representation centroid position and target centroid position. **b.** 100 channel Braincodec patterns for 10 x -by-10 LED array. 16 pure tones logarithmically spaced from 500 to 2000 Hz were passed to the trained encoder to produce 16, 10 x ×10 10×10 Braincodec spatiotemporal patterns. **c.** Time-averaged 10 x ×1010×10 stimulation patterns for each stimulus.

**Extended Data Fig. 7:**
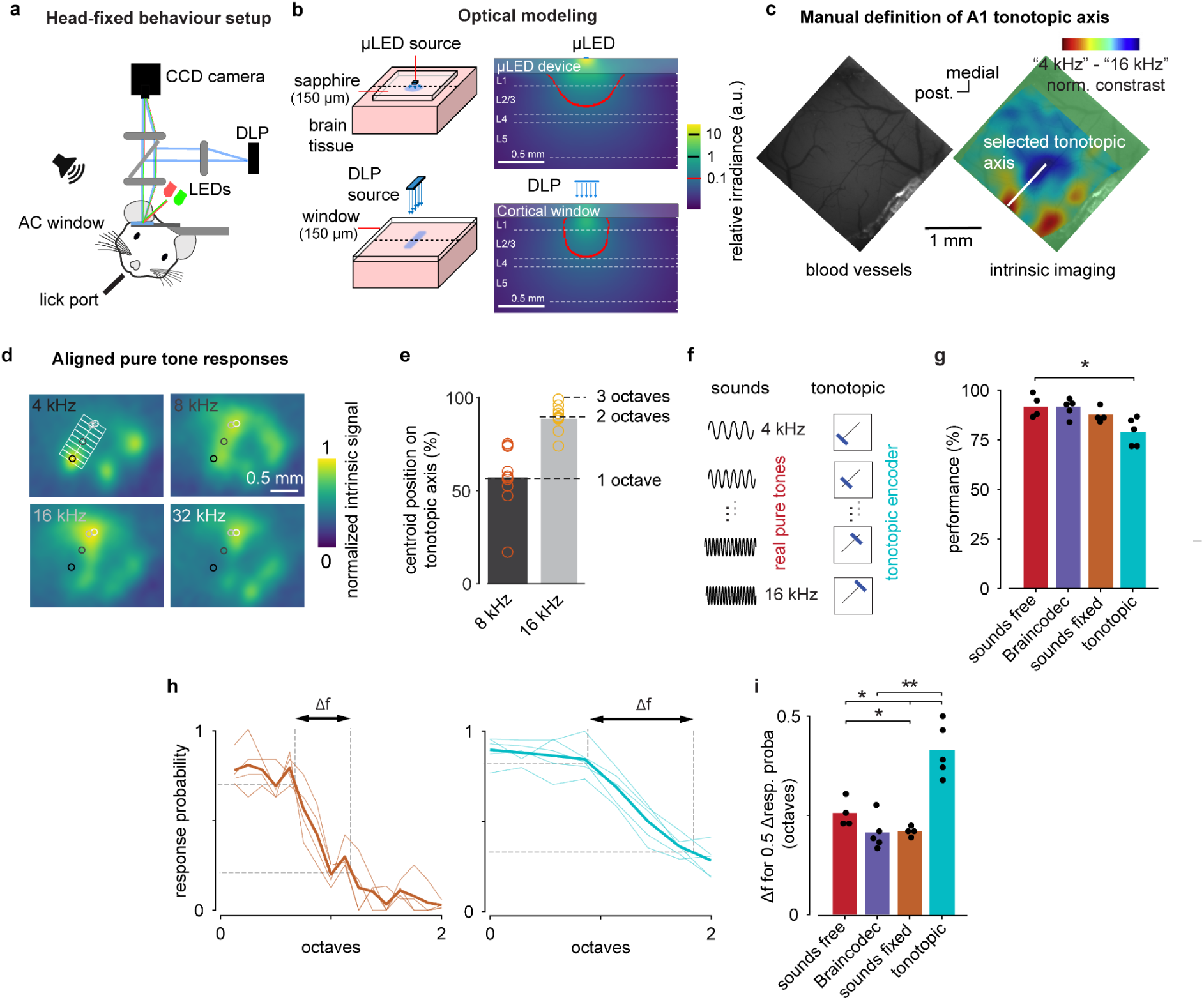
Behavioral task settings and tonotopic map alignment. **a.** Sketch of the head-fixed setup for stimulus discrimination training. The auditory cortex (AC) is stimulated with light delivered by a Digital Light Processor (DLP) through a cranial window. The same setup is used for intrinsic imaging (red LED) or blood vessel pattern imaging (green LED) with a CCD camera. **b.** (left) Schematic of the µLED (top) and DLP (bottom) light diffusion models (Zemax OpticStudio 2021). The dashed line indicates the cross-section that was taken for the optical profiles. (right) Optical profiles showing light delivery through brain tissue with a µLED (top) and with the DLP (bottom). Irradiance is normalised to be equal to 1 at the cortical surface in both cases, and the contour indicates the 10% level. The simulation indicates that the spatial resolution of light delivery in the cortex with the DLP is higher than the µLED array, as the DLP provides a smaller light cone angle at cortical surface, but the difference is limited due to the strong influence of light scattering in the tissue. **c.** (left) In the head-fixed setup, optogenetic stimuli are reproducibly positioned in every experimental session thanks to the stable landmarks provided by the blood vessel patterns. (right) The tonotopic axis of A1 was identified within these landmarks from intrinsic imaging maps obtained under anesthesia, before the behavioral training, as the axis joining the most posterior response maximum for a 4 kHz tone, and the maximum of the response to a 16 kHz tone. **d**. The tonotopic model consists in encoding a particular pure tone frequency as a 125 x 560 µm bar perpendicular to the A1 tonotopic axis. Eight non overlapping bars represented pure tones from low to high frequency. To estimate the correspondence between stimulation area and frequency, we performed intrinsic imaging. Intrinsic imaging responses to 4, 8, 16 and 32 kHz tone averaged across 10 C57Bl6J mice are superimposed with the centroids of the posterior part of the responses (black to light grey circle = 4 to 32 kHz). The axis joining these centroids delineates A1. The rectangles superimposed to the response map are the locations of the stimulus bars. The response maps indicate that 8 kHz response overlaps with the 4 and 16 kHz domains, but has a centroid lying in between them, and that the 16 kHz and 32 kHz domains in A1 are highly overlapping with centroids very close to each other (see **e**). Therefore, the low frequency in the tonotopic model corresponds to ∼4kHz, and high frequency to 16 to 32kHz. For the estimation of the resolution of perceived frequencies we chose to assign 16 kHz to the high frequency region to cover 2 octaves as in the pure tone discrimination experiment. Due to the similarity of 16 and 32 kHz responses, this choice likely overestimates the resolution of perception. **e.** Quantification of the distance between the centroid of the 4 kHz response in A1 and the centroid of the 8 and 16 kHz response, normalized by the distance between the centroids of the 4 kHz and 32 kHz response measured individually in each of the 10 mice shown in **d**. The centroids of 16 and 32 kHz are within 10% (∼100µm) of the full extent of the tonotopic axis (4 to 32kHz). **f.** Sketch of the categorization tasks for each group of mice. The “tonotopic” group discriminates between the two optogenetic stimulation bars located at each extreme of the tonotopic axis in a head-fixed Go/Nogo task. The “sound fixed” group discriminated between 4 and 16 kHz pure tones under head-fixation. After learning, mice of each group were presented at low probability with intermediate stimuli depicted in the panel (spontaneous categorization task). **g.** Behavioral performance during the spontaneous categorization task (mean accuracy of Go and Nogo trials excluding catch trials) for the groups performing ‘Braincodec’ or ‘tonotopic’ pattern discrimination or sound discrimination in the freely moving (‘sounds free’) or head-fixed (‘sounds fixed’) setting. The ‘Braincodec’ and ‘sound free’ conditions are shown in Fig. 5. Two-sided Wilcoxon rank sum tests, Braincodec vs tonotopic, p=0.032; sounds free vs fixed: p = 0.11. **h.** Categorization curves for the ‘tonotopic’ stimulation (blue) and for sounds in head-fixation (orange). Thin lines: single animals. Thick line: group average. Dashed lines: sketch the perceptual resolution measure. **i**. Perceptual resolution measured as the minimal frequency difference for which response probabilities differ by 0.5. Two-sided Wilcoxon rank sum tests. Braincodec vs tonotopic, p = 0.008; tonotopic vs sound free & fixed, p = 0.016 & 0.016; sound free vs fixed, p = 0.029; Sounds free: n = 4, Braincodec: n = 5; sounds fixed: n = 4, tonotopic: n = 5 mice; dots = single mice; bar = group average).

## Supplementary information

**Supplementary Video 1: SI-SNR for evaluating audio quality.** This video presents a 1 s music sample and five degraded versions of the same sample to illustrate the relationship between sound degradation and SI-SNR. Spectrograms are displayed simultaneously with the corresponding audio samples. The SI-SNR value and degradation level of each sample are displayed before playback.

**Supplementary Video 2: Braincodec reconstruction examples.** This video presents five original sound samples (speech, music, frog vocalization, washing machine) each followed by their reconstruction obtained with a deep neural network decoder after encoding by the 512 channel Braincodec algorithm. Spectrograms are displayed simultaneously with the corresponding audio samples.

**Supplementary Video 3: Comparison of cochlear implant synthesis and Braincodec for a speech sound.** This video presents a speech sound sample followed by 1) its reconstruction by a deep decoder after encoding by the 512 channel Braincodec algorithm, 2) its reconstruction by a vanilla vocoder (traditional approach) after the first processing step of the 22 channel ACE encoder of a cochlear implant, 3) same as 2) but reconstruction is done with a deep neural network decoder, 4) same as 3) but after the last encoding step of the ACE encoder. Spectrograms are displayed simultaneously with the corresponding audio samples.

**Supplementary Video 4: Comparison of cochlear implant synthesis and Braincodec for a music sound.** This video presents a music sound sample followed by 1) its reconstruction by a deep decoder after encoding by the 512 channel Braincodec algorithm, 2) its reconstruction by a vanilla vocoder (traditional approach) after the first processing step of the 22 channel ACE encoder of a cochlear implant, 3) same as 2) but reconstruction is done with a deep neural network decoder, 4) same as 3) but after the last encoding step of the ACE encoder. Spectrograms are displayed simultaneously with the corresponding audio samples.

**Supplementary Video 5: Comparison of cochlear implant synthesis and Braincodec for a natural sound.** This video presents a frog vocalization sound sample followed by 1) its reconstruction by a deep decoder after encoding by the 512 channel Braincodec algorithm, 2) its reconstruction by a vanilla vocoder (traditional approach) after the first processing step of the 22 channel ACE encoder of a cochlear implant, 3) same as 2) but reconstruction is done with a deep neural network decoder, 4) same as 3) but after the last encoding step of the ACE encoder. Spectrograms are displayed simultaneously with the corresponding audio samples.

**Supplementary Video 6: Comparison of cochlear implant synthesis and Braincodec for a machine sound.** This video presents a washing machine sound sample followed by 1) its reconstruction by a deep decoder after encoding by the 512 channel Braincodec algorithm, 2) its reconstruction by a vanilla vocoder (traditional approach) after the first processing step of the 22 channel ACE encoder of a cochlear implant, 3) same as 2) but reconstruction is done with a deep neural network decoder, 4) same as 3) but after the last encoding step of the ACE encoder. Spectrograms are displayed simultaneously with the corresponding audio samples.

**Supplementary Audio 1: Stimuli to probe channel tuning to auditory features.** 1 s audio samples of the stimuli used in **Fig. 3b**. Pure tones (PT) at 250 Hz, 1 kHz, 4 kHz to probe frequency tuning. Chirps from 250 Hz to 1 kHz and from 1 kHz to 250 Hz to probe frequency modulation (FM) tuning. Amplitude modulations (AM) with a carrier frequency of 1 kHz and modulations at 100, 30, 10 and 4 Hz to probe amplitude modulation tuning. Complex sounds of speech, music and natural sound (frog).

**Supplementary Audio 2: CIS cochlear--implant synthesis sounds.** Audio samples of various sounds and their reconstruction using a 12--channel CIS cochlear-implant strategy.

**Supplementary Audio 3: Stimuli to probe channel tuning to sound localisation.** 1 s audio samples of the stimuli used in **Fig. 6e** for the stereo variant of 512 channels Braincodec. Pure tones (PT) at 0.5 kHz coming from the left, front and right sides. Amplitude modulations (AM) with a carrier frequency of 1 kHz and modulations at 4.5 Hz coming from the left, front and right sides. Music sound sample coming from the left, front and right sides.

## References

1. Zeng, F.-G., Qi, J., Wu, C.-C., Shu, Y. & Chai, R. Treating Hearing Loss: From Cochlear Implantation to Gene Therapy. Adv. Sci. 12, e09960 (2025).

2. Macherey, O. & Carlyon, R. P. Cochlear implants. Curr. Biol. CB 24, R878–R884 (2014).

3. Hamilton, L. S., Oganian, Y., Hall, J. & Chang, E. F. Parallel and distributed encoding of speech across human auditory cortex. Cell 184, 4626–4639.e13 (2021).

4. Penfield, W. & Perot, P. The Brain’s Record of Auditory and Visual Experience. A final summary and discussion. Brain J. Neurol. 86, 595–696 (1963).

5. Zeng, F.-G. Celebrating the one millionth cochlear implant. JASA Express Lett. 2, 077201 (2022).

6. Trouillet, A. et al. High-resolution prosthetic hearing with a soft auditory brainstem implant in macaques. Nat. Biomed. Eng. 1–15 (2025) doi:10.1038/s41551-025-01378-9.

7. Wong, K. et al. Auditory Brainstem Implants: Recent Progress and Future Perspectives. Front. Neurosci. 13, 10 (2019).

8. Beauchamp, M. S. et al. Dynamic Stimulation of Visual Cortex Produces Form Vision in Sighted and Blind Humans. Cell 181, 774–783.e5 (2020).

9. Palanker, D., Weiland, J. D., Rosin, B. & Sahel, J.-A. Restoration of Sight with Electronic Retinal Prostheses. Annu. Rev. Vis. Sci. 11, 125–147 (2025).

10. Fernández, E. et al. Visual percepts evoked with an intracortical 96-channel microelectrode array inserted in human occipital cortex. J. Clin. Invest. 131, e151331 (2021).

11. Sahel, J.-A. et al. Partial recovery of visual function in a blind patient after optogenetic therapy. Nat. Med. 27, 1223–1229 (2021).

12. Valle, G. et al. Tactile edges and motion via patterned microstimulation of the human somatosensory cortex. Science 387, 315–322 (2025).

13. Nl, D. & Jt, R. Auditory Brainstem Implantation: Candidacy Evaluation, Operative Technique, and Outcomes. Otolaryngol. Clin. North Am. 53, (2020).

14. Wagner, L., Altindal, R., Plontke, S. K. & Rahne, T. Pure tone discrimination with cochlear implants and filter-band spread. Sci. Rep. 11, 20236 (2021).

15. Chen, X., Wang, F., Fernandez, E. & Roelfsema, P. R. Shape perception via a high-channel-count neuroprosthesis in monkey visual cortex. Science 370, 1191–1196 (2020).

16. Taylor, J. A. et al. Cortical neuroprostheses improve auditory coding and perception compared to cochlear implants. 2025.10.27.684780 Preprint at 10.1101/2025.10.27.684780 (2025).

17. Uguz, I. & Shepard, K. L. Spatially controlled, bipolar, cortical stimulation with high-capacitance, mechanically flexible subdural surface microelectrode arrays. Sci. Adv. 8, eabq6354 (2022).

18. Greer, R. et al. Chronically implantable μLED arrays for optogenetic cortical surface stimulation in mice. Nat. Commun. 17, 878 (2026).

19. Heil, P. & Peterson, A. J. Basic response properties of auditory nerve fibers: a review. Cell Tissue Res. 361, 129–158 (2015).

20. Wang, X., Lu, T., Bendor, D. & Bartlett, E. Neural coding of temporal information in auditory thalamus and cortex. Neuroscience 154, 294–303 (2008).

21. Koumura, T., Terashima, H. & Furukawa, S. Cascaded Tuning to Amplitude Modulation for Natural Sound Recognition. J. Neurosci. 39, 5517–5533 (2019).

22. Dick, F. K. et al. Extensive Tonotopic Mapping across Auditory Cortex Is Recapitulated by Spectrally Directed Attention and Systematically Related to Cortical Myeloarchitecture. J. Neurosci. 37, 12187–12201 (2017).

23. Hackett, T. A., Preuss, T. M. & Kaas, J. H. Architectonic identification of the core region in auditory cortex of macaques, chimpanzees, and humans. J. Comp. Neurol. 441, 197–222 (2001).

24. Norman-Haignere, S. V. et al. Temporal integration in human auditory cortex is predominantly yoked to absolute time. Nat. Neurosci. 28, 2356–2365 (2025).

25. Osman, A. F., Lee, C. M., Escabí, M. A. & Read, H. L. A Hierarchy of Time Scales for Discriminating and Classifying the Temporal Shape of Sound in Three Auditory Cortical Fields. J. Neurosci. 38, 6967–6982 (2018).

26. Kopp-Scheinpflug, C., Sinclair, J. L. & Linden, J. F. When Sound Stops: Offset Responses in the Auditory System. Trends Neurosci. 41, 712–728 (2018).

27. Lamothe, C., Bagur, S., Gosselin, E. & Bathellier, B. Sound offset responses become highly informative in the auditory cortex. J. Physiol. https://doi.org/10.1113/JP288416 (2025) doi:10.1113/JP288416.

28. Bagur, S. et al. A spatial code for temporal information is necessary for efficient sensory learning. Sci. Adv. 11, eadr6214 (2025).

29. Buck, A. et al. Neural Response Reliability as a Marker of the Transition of Neural Codes along Auditory Pathways. Adv. Sci. 12, e08777 (2025).

30. Wang, X. Cortical Coding of Auditory Features. Annu. Rev. Neurosci. 41, 527–552 (2018).

31. Gosselin, E. et al. Sound feature representations decorrelate across the mouse auditory pathway. PLOS Biol. 23, e3003452 (2025).

32. Kell, A. J. E., Yamins, D. L. K., Shook, E. N., Norman-Haignere, S. V. & McDermott, J. H. A Task-Optimized Neural Network Replicates Human Auditory Behavior, Predicts Brain Responses, and Reveals a Cortical Processing Hierarchy. Neuron 98, 630–644.e16 (2018).

33. Tuckute, G., Feather, J., Boebinger, D. & McDermott, J. H. Many but not all deep neural network audio models capture brain responses and exhibit correspondence between model stages and brain regions. PLoS Biol. 21, e3002366 (2023).

34. Défossez, A., Copet, J., Synnaeve, G. & Adi, Y. High Fidelity Neural Audio Compression. Preprint at 10.48550/arXiv.2210.13438 (2022).

35. Bellier, L. et al. Music can be reconstructed from human auditory cortex activity using nonlinear decoding models. PLOS Biol. 21, e3002176 (2023).

36. Pasley, B. N. et al. Reconstructing speech from human auditory cortex. PLoS Biol 10, e1001251 (2012).

37. Krumbholz, K., Patterson, R. D. & Pressnitzer, D. The lower limit of pitch as determined by rate discrimination. J. Acoust. Soc. Am. 108, 1170–1180 (2000).

38. Zeghidour, N., Luebs, A., Omran, A., Skoglund, J. & Tagliasacchi, M. SoundStream: An End-to-End Neural Audio Codec. Preprint at 10.48550/arXiv.2107.03312 (2021).

39. Hinton, G. E. & Salakhutdinov, R. R. Reducing the dimensionality of data with neural networks. Science 313, 504–7 (2006).

40. Conwell, C., Prince, J. S., Kay, K. N., Alvarez, G. A. & Konkle, T. A large-scale examination of inductive biases shaping high-level visual representation in brains and machines. Nat. Commun. 15, 9383 (2024).

41. Yamins, D. L. K. & DiCarlo, J. J. Using goal-driven deep learning models to understand sensory cortex. Nat. Neurosci. 19, 356–365 (2016).

42. Luo, Y. & Mesgarani, N. TasNet: time-domain audio separation network for real-time, single-channel speech separation. Preprint at 10.48550/arXiv.1711.00541 (2018).

43. Norman-Haignere, S., Kanwisher, N. G. & McDermott, J. H. Distinct Cortical Pathways for Music and Speech Revealed by Hypothesis-Free Voxel Decomposition. Neuron 88, 1281–1296 (2015).

44. Gaucher, Q. et al. Complexity of frequency receptive fields predicts tonotopic variability across species. eLife 9, e53462 (2020).

45. Leonard, M. K. et al. Large-scale single-neuron speech sound encoding across the depth of human cortex. Nature 1–10 (2023) doi:10.1038/s41586-023-06839-2.

46. Deneux, T., Kempf, A., Daret, A., Ponsot, E. & Bathellier, B. Temporal asymmetries in auditory coding and perception reflect multi-layered nonlinearities. Nat Commun 7, 12682 (2016).

47. Aponte, D. A. et al. Recurrent network dynamics shape direction selectivity in primary auditory cortex. Nat. Commun. 12, 314 (2021).

48. Bhumika, S. et al. A Late Critical Period for Frequency Modulated Sweeps in the Mouse Auditory System. Cereb. Cortex N. Y. N 1991 30, 2586–2599 (2020).

49. Altmann, C. F. & Gaese, B. H. Representation of frequency-modulated sounds in the human brain. Hear. Res. 307, 74–85 (2014).

50. Vandali, A. E., Whitford, L. A., Plant, K. L. & Clark, G. M. Speech perception as a function of electrical stimulation rate: using the Nucleus 24 cochlear implant system. Ear Hear. 21, 608–624 (2000).

51. Karoui, C. et al. Searching for the Sound of a Cochlear Implant: Evaluation of Different Vocoder Parameters by Cochlear Implant Users With Single-Sided Deafness. Trends Hear. 23, 2331216519866029 (2019).

52. de Nobel, J., Briaire, J. J., Bäck, T. H. W., Kononova, A. V. & Frijns, J. H. M. From spikes to speech: NeuroVoc — A biologically plausible vocoder framework for auditory perception and cochlear implant simulation. Hear. Res. 469, 109469 (2026).

53. Drinnenberg, A. et al. Large-scale cellular-resolution read/write of activity enables discovery of cell types defined by complex circuit properties. 2025.10.21.683734 Preprint at 10.1101/2025.10.21.683734 (2025).

54. Theunissen, F. E. & Elie, J. E. Neural processing of natural sounds. Nat. Rev. Neurosci. 15, 355–366 (2014).

55. Wood, K. C., Town, S. M. & Bizley, J. K. Neurons in primary auditory cortex represent sound source location in a cue-invariant manner. Nat. Commun. 10, 3019 (2019).

56. Saddler, M. R. & McDermott, J. H. Models optimized for real-world tasks reveal the task-dependent necessity of precise temporal coding in hearing. Nat. Commun. 15, 10590 (2024).

57. Mrsic-Flogel, T. D., King, A. J. & Schnupp, J. W. H. Encoding of virtual acoustic space stimuli by neurons in ferret primary auditory cortex. J. Neurophysiol. 93, 3489–3503 (2005).

58. Mathis, M. W., Perez Rotondo, A., Chang, E. F., Tolias, A. S. & Mathis, A. Decoding the brain: From neural representations to mechanistic models. Cell 187, 5814–5832 (2024).

59. Drakopoulos, F. et al. Modelling neural coding in the auditory midbrain with high resolution and accuracy. Nat. Mach. Intell. 7, 1478–1493 (2025).

60. Wingert, J. C., Parida, S., Norman-Haignere, S. V. & David, S. V. Convolutional neural network models describe the encoding subspace of local circuits in auditory cortex. Nat. Neurosci. 29, 876–887 (2026).

61. Leonard, M. K. et al. Large-scale single-neuron speech sound encoding across the depth of human cortex. Nature 626, 593–602 (2024).

62. Hettick, M. et al. Minimally invasive implantation of scalable high-density cortical microelectrode arrays for multimodal neural decoding and stimulation. Nat. Biomed. Eng. 10, 1206–1221 (2026).

63. Greenspon, C. M. et al. Long-term safety and efficacy of intracortical microstimulation in humans. Sci. Transl. Med. 18, eaec3728 (2026).

64. Valle, G. et al. Restoring vision and touch with cortical microstimulation. Nat. Rev. Bioeng. 1–20 (2026) doi:10.1038/s44222-026-00449-z.

65. Stronks, H. C. et al. Effects of Preoperative Factors on the Learning Curves of Postlingual Cochlear Implant Recipients. Ear Hear. 46, 1560–1572 (2025).

66. Paszke, A., et al. PyTorch: An Imperative Style, High-Performance Deep Learning Library. Preprint at 10.48550/arXiv.1912.01703 (2019).

67. Kingma, D. P. & Ba, J. Adam: A Method for Stochastic Optimization. Preprint at 10.48550/arXiv.1412.6980 (2017).

68. Feather, J., Durango, A., Gonzalez, R. & McDermott, J. Metamers of neural networks reveal divergence from human perceptual systems. in Advances in Neural Information Processing Systems vol. 32 (Curran Associates, Inc., 2019).

69. Baumann, T., Köhn, A. & Hennig, F. The Spoken Wikipedia Corpus collection: Harvesting, alignment and an application to hyperlistening. Lang. Resour. Eval. 53, 303–329 (2019).

70. Hershey, S., et al. The Benefit Of Temporally-Strong Labels In Audio Event Classification. Preprint at 10.48550/arXiv.2105.07031 (2021).

71. Gemmeke, J. F. et al. Audio Set: An ontology and human-labeled dataset for audio events. in 2017 IEEE International Conference on Acoustics, Speech and Signal Processing (ICASSP) 776–780 (2017). doi:10.1109/ICASSP.2017.7952261.

72. Piczak, K. J. ESC: Dataset for Environmental Sound Classification. in Proceedings of the 23rd ACM international conference on Multimedia 1015–1018 (Association for Computing Machinery, New York, NY, USA, 2015). doi:10.1145/2733373.2806390.

73. Julich-Brain: A 3D probabilistic atlas of the human brain’s cytoarchitecture. Science https://www.science.org/doi/10.1126/science.abb4588.

74. Spearman, C. The Proof and Measurement of Association between Two Things. Am. J. Psychol. 15, 72–101 (1904).

75. Huth, A. G., de Heer, W. A., Griffiths, T. L., Theunissen, F. E. & Gallant, J. L. Natural speech reveals the semantic maps that tile human cerebral cortex. Nature 532, 453–458 (2016).

76. Nucleus Toolbox. https://fr.mathworks.com/matlabcentral/fileexchange/159688-nucleus-toolbox (2026).

77. Verdier, A., Aldanondo, A., Bathellier, B. & Bagur, S. Enhanced performance in perceptual decision making task without deprivation in mice using medial forebrain bundle stimulation. 2022.03.29.486230 Preprint at 10.1101/2022.03.29.486230 (2022).

78. Ceballo, S., Piwkowska, Z., Bourg, J., Daret, A. & Bathellier, B. Targeted Cortical Manipulation of Auditory Perception. Neuron 104, 1168–1179.e5 (2019).

79. Ceballo, S. et al. Cortical recruitment determines learning dynamics and strategy. Nat Commun 10, 1479 (2019).

